# Nitric oxide, K_ATP_ channels and endothelin-1 modulate brain pericyte function, vascular tone and neurovascular coupling

**DOI:** 10.1101/2020.06.07.138875

**Authors:** Stefan Andreas Zambach, Changsi Cai, Hans Christian Cederberg Helms, Bjørn Olav Hald, Jonas Christoffer Fordsmann, Reena Murmu Nielsen, Micael Lønstrup, Birger Brodin, Martin Johannes Lauritzen

## Abstract

Neurotransmitter-mediated signaling correlates strongly to changes in cerebral blood flow (CBF), and functional neuroimaging relies on the robust coupling between activity and CBF, i.e. neurovascular coupling (NVC). We here reveal that key endothelial signaling molecules, nitric oxide (eNO) and endothelin-1 (ET1), modulate pericyte contractility and that pericyte ATP-sensitive potassium (K_ATP_) channels interact with endothelial factors to modulate vascular tone and NVC. We show that NVC requires local synthesis of cGMP, but not NO derived from endothelial cells. The potent endothelial vasoconstrictor ET1 contracted pericytes by IP_3_ receptor mediated Ca^2+^ release and blocked NVC. In comparison, pericyte K_ATP_ channel openers increased the diameter of capillaries by deactivation of L-type Ca^2+^ channels while K_ATP_ blockers shortened the NVC response. All vasoactive stimuli produced the largest diameter changes at the first capillary that branches off from the penetrating arteriole. Our results reveal that three different signaling pathways mediate the effects of NO, ET1 and K_ATP_ channels on brain pericytes and capillary blood flow by mechanisms similar to vascular smooth muscle despite great differences in morphology.

## INTRODUCTION

Variations in neuronal activity are accompanied by robust changes in brain metabolism and perfusion. This relationship is exploited by functional neuroimaging techniques that use changes in cerebral blood flow (CBF) or the blood-oxygenation level-dependent signal to track brain function [1]. The structural basis for local blood flow control is the neurovascular unit (NVU), which comprises neurons, astrocytes, vascular mural cells, endothelial cells and components of the extracellular matrix [2-4]. The term “neurovascular coupling” (NVC) denotes the robust regulation of CBF that accompany changes in brain activity, in order to match energy demands. Disruption of NVC is a key characteristic of neurologic diseases, e.g. Alzheimer’s disease [5], stroke [6] and sub-arachnoid hemorrhage [7]. Therefore, there is an urgent need to understand NVC mechanisms.

Vascular smooth muscle cells (VSMCs) encircle brain arteries and arterioles, representing a key mechanism of cerebrovascular resistance, and pericytes on adjacent capillaries may serve a similar regulatory function [8, 9]. Novel findings suggest that compared to VSMCs on arteries and arterioles, precapillary sphincters and pericytes on the first few branches of capillaries produce larger and faster vascular diameter responses to any type of stimulation [8-10]. Furthermore, endothelial pathways are involved in NVC [11], possibly via NMDA receptors on capillary endothelial cells [12] or Caveolin-1 on arteriolar endothelial cells [4]. However, how signaling molecules from brain endothelial cells (BECs) may influence pericytes and contribute to capillary blood flow is incompletely understood [4].

Endothelium-derived factors such as the dilator nitric oxide (NO) and the constrictor endothelin-1 (ET1) modulate vascular tone in arteries and arterioles, but how BECs influence pericytes and thus brain capillaries is unclear. Our first objective was to examine the involvement of the NO and ET1 signaling cascade in pericytes and blood flow regulation in brain capillaries in the resting state, during activity increases and in cardiac arrest. Our second objective was to identify the pericyte signaling pathways involved in NO-mediated relaxation and ET1-induced constriction by using isolated pericytes. Because NOS inhibitors may modulate ATP-sensitive potassium (K_ATP_) channels, which are highly enriched in pericytes, and contribute to capillary flow regulation and NVC, our third objective was to exclude a role for K_ATP_ channels in the observed effects of NO and to explore the role of K_ATP_ channels in pericytes and capillary blood flow regulation. Our findings provide a mechanistic insight into endothelium-derived signaling NO and ET1 in diameter modulation of penetrating arterioles (PAs) and their associated capillaries, as well as in modulation of NVC, but we provide evidence that the effect of K_ATP_ channels is independent of these processes. Pericytes at 1^st^ -order capillaries are vascular hubs for NVC, and modifications in cerebrovascular resistance may depend on this vascular segment.

## RESULTS

We used *in vivo* two-photon microscopy to examine the regulation of capillary blood flow and NVC in anesthetized NG2dsRed mice, which express a red fluorescent protein in pericytes (in red, **Fig. 1C**) and in VSMCs [13]. We identified capillaries by their branching order, with 1 order being the first capillary branching from the PA, and so on (**Fig. 1C**). To minimize focus drift of *in vivo* two-photon microscopy, we implemented fast and repetitive three-dimensional (3D) imaging, with 10 – 14 planes per image stack per second. A glass micropipette was inserted in proximity to the PA and the 1 – 3 order capillaries for local pressure-ejection (puff) of vasoactive compounds and for recording of local field potentials (LFPs) (**Fig. 1A – C**) [9, 14]. Whisker pad (WP) stimulation was used for NVC studies [8-10].

**Figure 1.**
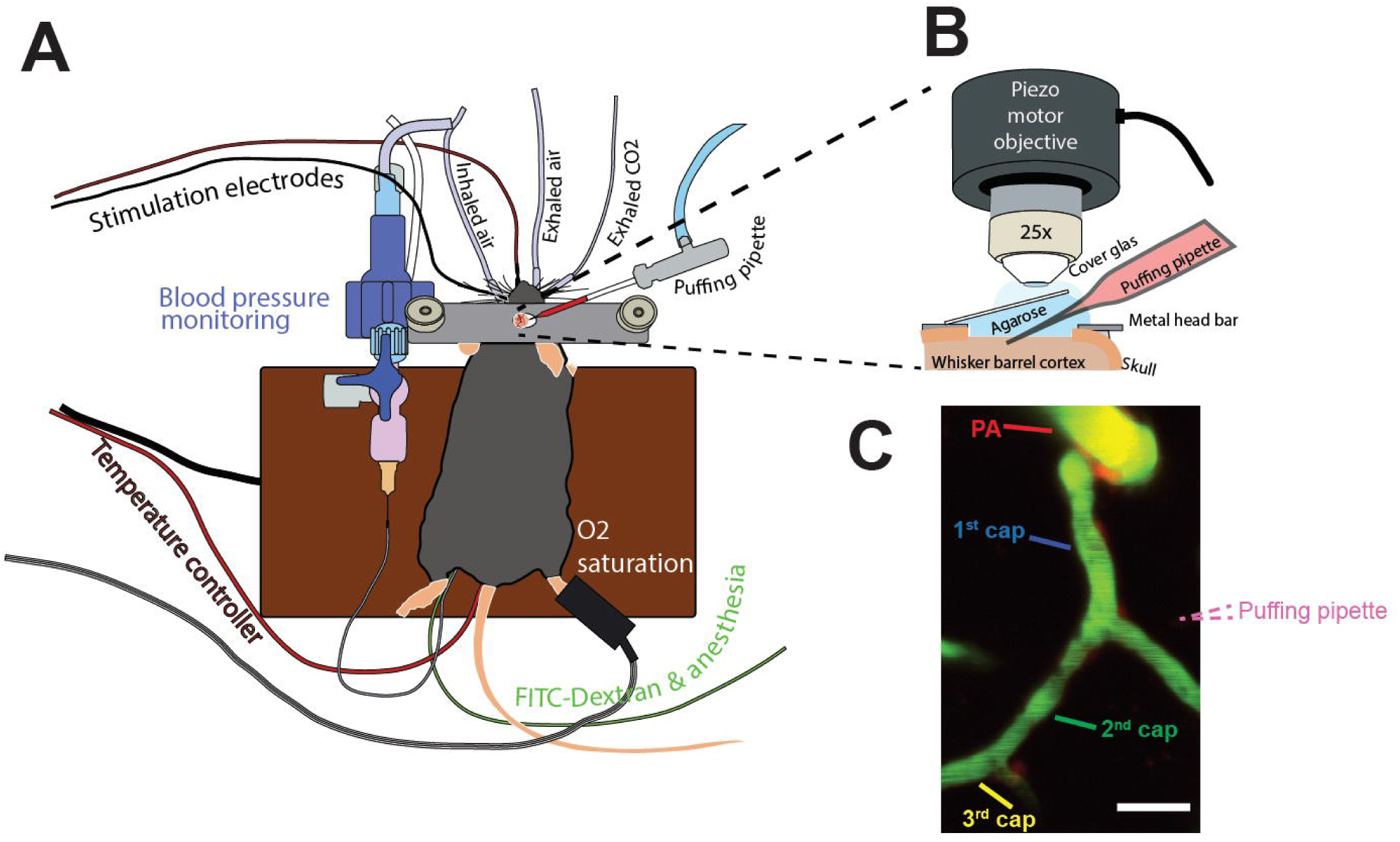
Diagram of the *in vivo* experimental setup. (**A**) The physiological state of the mouse was monitored throughout the experiment, including arterial blood pressure, exhaled CO_2_, body temperature and O_2_ saturation. Vessel lumen dye FITC-dextran and anesthesia alpha-chloralose were infused via a femoral vein catheter. Two stimulations are implemented: Whisker pad (WP) stimulation and a micro-glass pipette inserted into the imaging area for local ejection (puff). (**B**) The puffing micropipette is placed in proximity to the arteriole–capillary site, containing 10 μM Alexa 594 and vasoactive compound to visualize pipette tip as red color under the fluorescent microscope. The 25× objective with a piezo-motor is used to facilitate fast time-lapse 3D imaging. (**C**) An average of two-photon image stack recorded at different depths, with red indicating pericytes and the puffing pipette and green the vessel lumen. Capillaries are classified as the branching order from the penetrating arteriole (PA). The dashed red lines indicate the pipette tip. Scale bar: 20 µm.

To examine pericyte responses to vasodilatory and contractile signals, we imaged diameter changes of PAs and 1–3 order capillaries *in vivo* after delivery of vasodilatory and contractile pharmacological compounds. We monitored diameter rather than red blood cell flow; because flow measurements reflect events occurring in several segments of the microvascular bed in series, which conflate which segment or segments the mechanism affects [15]. In comparison, capillary diameter changes reflect mainly local responses, and if slowly propagating responses are involved, they can easily be recorded and considered [9].

### Acetylcholine (Ach)-induced vasodilation is mediated by NO synthesis in capillary endothelium

Pericytes engage in several types of surface contact with capillary BECs, including gap and adherens junctions and peg-socket junctions [16]. The peg-socket contacts involve reciprocal, interdigitating evaginations from each cell, which invaginate into each other. This proximity of endothelial cells and pericytes led us to examine the possibility of an endothelial influence on pericyte contractility. First, we examined the role of endothelial-derived NO for pericyte-mediated vasodilation. For this purpose, we recorded capillary diameter changes in response to local puffing of Ach, which activates the constitutive endothelial NO synthase (eNOS) causing NO release [17, 18]. Ach consistently induced vasodilation of the PA and the 1^st^ order capillaries, whereas an intravenous (i.v.) bolus infusion of the NOS inhibitor NG-nitro-L-arginine methyl ester (L-NAME) blocked the response (**Fig. 2A – C**). This result is consistent with endothelial NO as the main relaxing factor.

**Figure 2.**
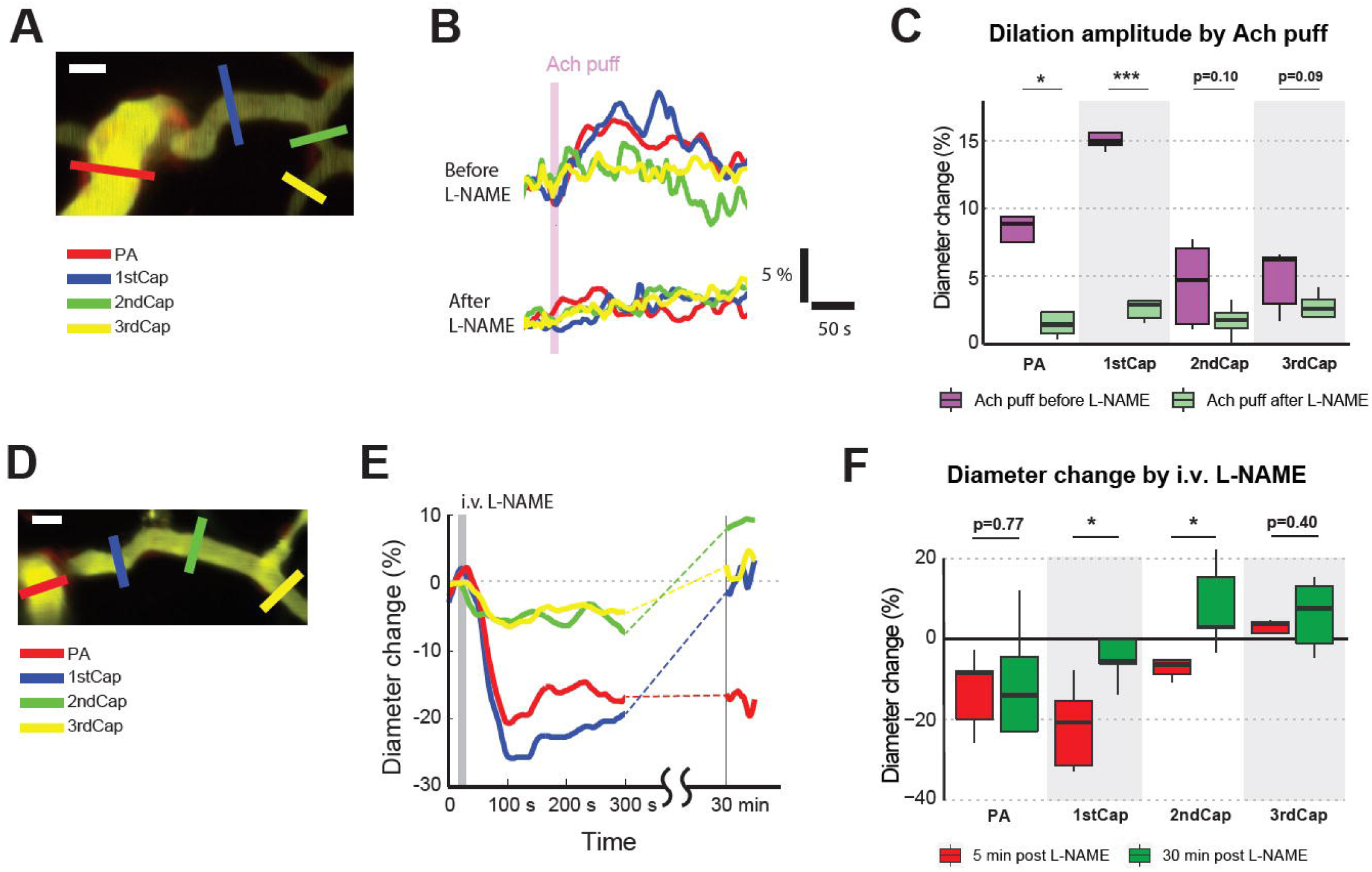
Acetylcholine (Ach)-induced vasodilation is mediated by endothelial nitric oxide synthesis (eNOS). (**A**) An average-projected image from a local two-photon image stack with color-coded regions of interest (ROIs) placed across PA and 1–3 order capillaries. (**B**) Diameter change of the vessel segment at each ROI was estimated (See Methods). Representative traces of Ach-puff-induced vasodilation before and after i.v. bolus infusion of L-NAME, a non-specific NOS inhibitor. The ROIs and the trace colors are coded identically. Each trace is normalized to its mean pre-puff value. (**C**) Comparison of relative diameter change at PA and 1–3 order capillaries, induced by Ach-puff before and after L-NAME. Before L-NAME: number of animal N=5, Number of vasculature n=10; after L-NAME: N=5, n=8. (**D**) An average-projected image from a local two-photon image stack with PA and 1–3 order capillaries. (**E**) Representative traces of vessel diameter change from onset of L-NAME infusion until 30 min post-infusion. (**F**) Comparison of relative diameter change at PA and 1–3 order capillaries at 5 min and 30 min post-infusion of L-NAME. After 30 min, PA remained constricted while 1–3 order capillaries returned to the pre-infusion state or even dilated. N=5, n=5. Paired or unpaired t-test, as appropriate. Scale bar: 10 µm.

To investigate whether endothelial NOS influence NVC, we used i.v. bolus injection of L-NAME and assessed the dilation peak, half-peak latency and half-peak duration evoked by WP stimulation. Systemic L-NAME affected none of the vascular responses, suggesting that eNOS activity increases are not crucial for a full development of NVC response (**Supplementary fig. 1**). L-NAME also had no effect on response amplitudes and latencies in WP-induced field excitatory post-synaptic potential (fEPSP) and field inhibitory post-synaptic potential (fIPSP) (**Supplementary fig. 2A – C**). However, it instantly reduced the baseline diameters of the PA and of 1–3 order capillaries, suggesting that constitutive eNOS activity accounts for a large portion of regulation in the basal vascular tone (**Fig. 2D – F**).

The arterial blood pressure increased immediately after i.v. administration of L-NAME and remained elevated for at least 30 minutes (**Table 1**). The rise in blood pressure represents systemic vasoconstriction from a lack of eNOS activity. PAs capillaries remained constricted at 30 minutes, whereas 1^st^, 2^nd^ and 3^rd^ order capillaries returned to the resting state at this time (**Fig. 2D – F**). This may imply that eNOS inhibition affects PA diameters persistently, but 1^st^ and higher orders capillaries are affected transiently.

**Table 1:** Arterial blood pressure before and after systemic administration of L-NAME

|  | Control | 5 min after<br>L-NAME | 30 min after<br>L-NAME |
| --- | --- | --- | --- |
| Blood<br>pressure<br>(mmHg) | 71 ± 6 | 112 ± 25 | 104 ± 21 |

### Vasodilation, induced by the NO donor SNAP, is mediated by cGMP in capillary pericytes

Results of *in vitro* experiments could suggest that VSMCs on PAs are sensitive to NO, but that pericytes on capillaries are not [19]. To address this question directly, we puffed the NO donor S-Nitroso-N-acetyl-DL-penicillamine (SNAP) onto the PA and onto 1**–**3 order capillaries and observed dilation of all segments with the largest response at the 1^st^ order capillary. We observed that both the PA and 1^st^ -3^rd^ order capillaries respond to NO (**Fig. 3A – C**). NO diffuses from endothelium into VSMCs and activates soluble guanylyl cyclase (sGC) to form cyclic guanosine monophosphate (cGMP), which triggers α-smooth muscle actin (SMA) relaxation [20, 21]. To examine whether sGC activity in pericytes mediates NO-induced capillary dilation, we locally puffed SNAP before and after topical application of the specific sGC inhibitor [1H-[1,2,4]oxadiazolo-[4, 3-a]quinoxalin-1-one] (ODQ). ODQ blunted the dilatory effect of SNAP puff (**Fig. 3A – C**), which supports the hypothesis that cGMP mediates NO-induced pericyte dilation.

**Figure 3.**
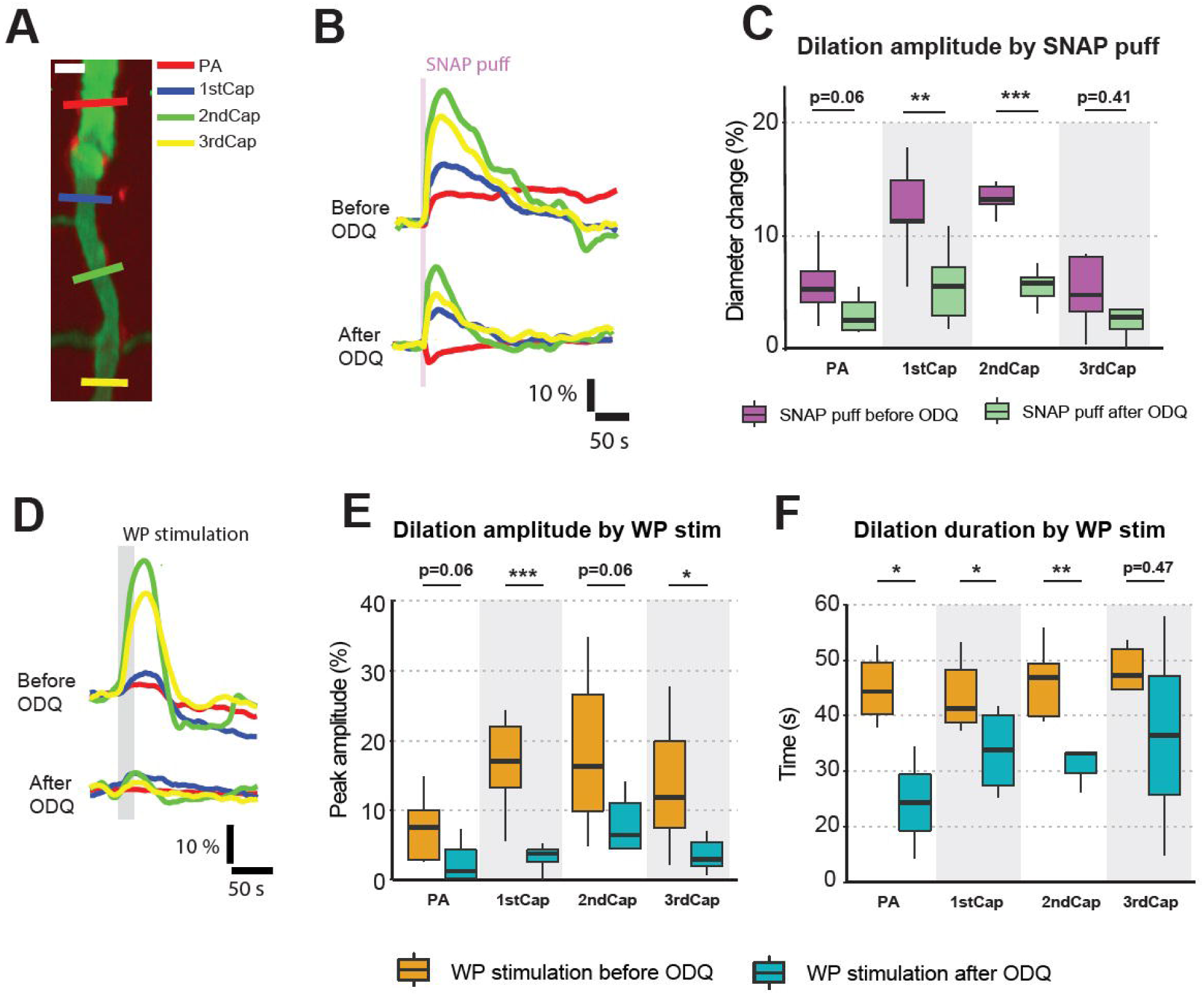
NO donor SNAP-induced vasodilation is mediated by cGMP in capillary pericytes. (A) An average-projected image from a local two-photon image stack with PA and 1–3 order capillaries. Scale bar: 10 µm. (**B**) Representative traces of SNAP-puff–induced vasodilation before and after topical application of cGMP inhibitor soluble guanylyl cyclase inhibitor (ODQ). (**C**) Comparison of relative diameter change at PA and 1–3 order capillaries, induced by SNAP-puff before and after ODQ. Before: N=7, n=7; after: N=6, n=6. (**D–F**) Comparison of WP stimulation induced vasodilation at PA and 1–3 order capillaries before and after ODQ, with (**D**) Representative traces of WP stimulation–induced vasodilation before and after topical application of ODQ. (**E**) Dilation amplitude. (**F**) Dilation duration of half-peak. Before: N=7, n=7; after: N=6, n=6. Paired or unpaired t-test, as appropriate.

Next, to examine whether cGMP participates in NVC, we compared WP-induced vasodilation before and after blocking sGC with ODQ. ODQ reduced the vasodilator response for all capillary orders and most profoundly for the 1^st^ order capillary (**Fig. 3D – E**). The half-peak duration was also shortened (**Fig. 3F**), whereas the half-peak latency to increase showed no significant change (**Supplementary fig. 3A**). In comparison, the evoked electrical responses, fEPSP and fIPSP, remained unchanged (**Supplementary fig. 2D – F**). These results imply that NO-dependent vasodilator responses in pericytes trigger increased cGMP, and that modulation of sGC activity contributes importantly to NVC.

### Pericyte K_ATP_ channels are closed in resting state and open in response to increased brain activity

NOS inhibitors may suppress the opening of K_ATP_ channels, which could complicates data interpretation [15, 22]. Furthermore, much existing knowledge about NO and K_ATP_ channels is based on studies using conductance vessels [23, 24], and whether NO and K_ATP_ channels interact in brain pericytes is unknown.

First, we assessed how local puffing of the K_ATP_ channel opener pinacidil affects pericytes. Pinacidil at 5 mM evoked dilation of PAs and capillaries with the same response amplitude as WP stimulation (**Fig. 4G**) and with the largest responses at the 1^st^ order capillary (**Fig. 4A – C**). This result suggested that K_ATP_ channels on VSMCs and pericytes of 1^st^ order capillaries have the potential to adjust the capillary diameter and blood flow according to metabolic needs. In comparison, the K_ATP_ channel closer PNU37883 (PNU) or sham cerebrospinal fluid (CSF) caused no significant vessel diameter change, suggesting that vascular K_ATP_ channels are closed in the resting state (**Fig. 4C, G**). Topical application of 0.5 mM PNU abolished the vasodilator effect of pinacidil (**Fig. 4A – C**), consistent with specificity of this vasodilator effect to K_ATP_ channels.

**Figure 4.**
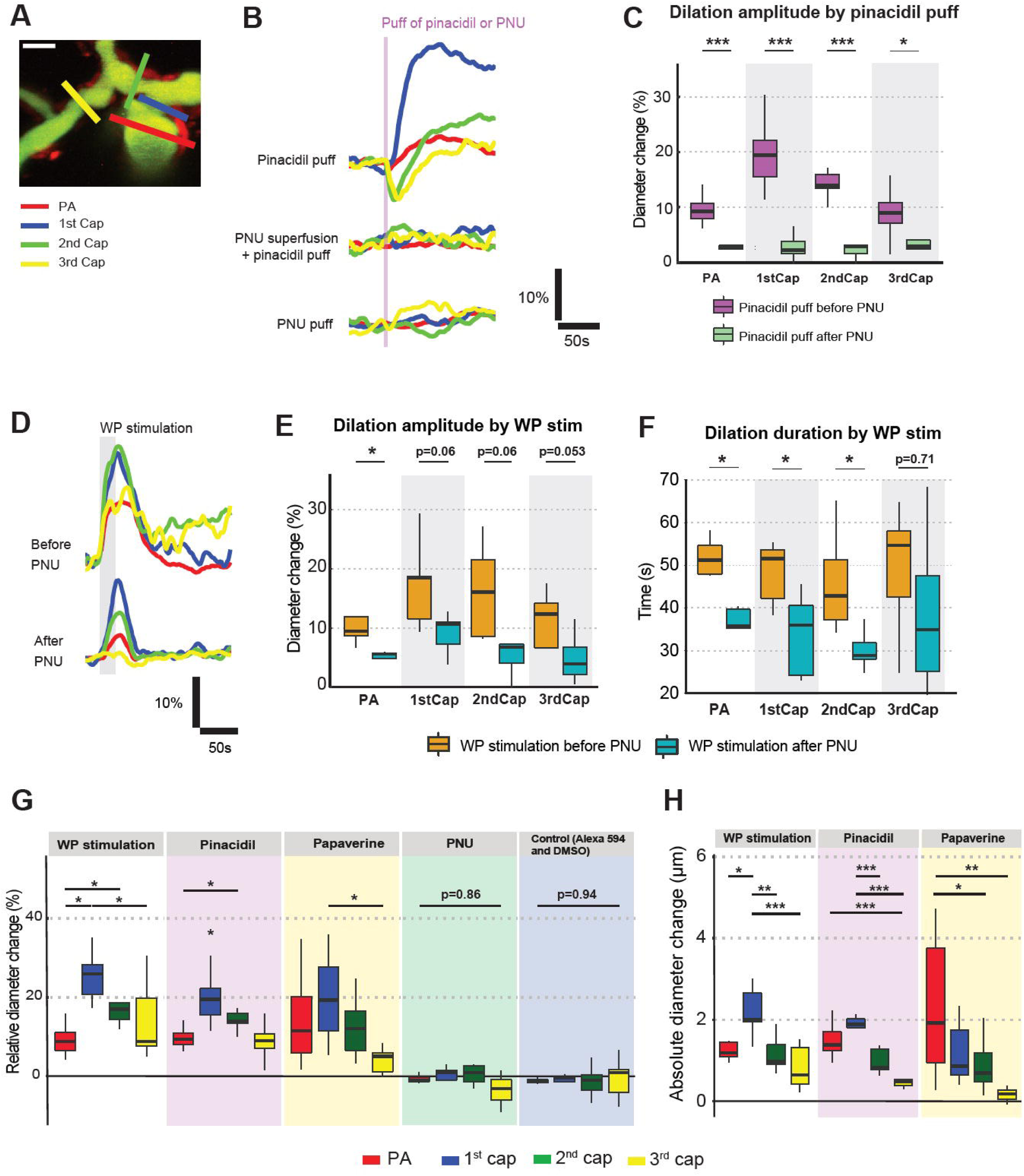
ATP-sensitive potassium (K_ATP_) channels enhance and prolong WP stimulation induced vasodilation. (**A**) An average-projected image from a local two-photon image stack with four color-coded ROIs placed across at PA, 1–3 order capillaries, respectively. Scale bar: 10 µm. (**B**) Representative traces of K_ATP_ channel opener pinacidil puff induced vasodilation before and after K_ATP_ channel closer PNU37883 (PNU), as well as a trace of vascular diameter following PNU puff. (**C**) Comparison of peak diameter change in percentage. Paired or unpaired t-test, as appropriate. Before: N=9, n=15; after: N=5, n=8. (**D**) Representative traces of WP stimulation induced vasodilation before and after topical application of PNU. (**E–F**) Comparison of vasodilation curves before and after topical application of PNU. (**E**) Peak diameter change in percentage. Before: N=5, n=14; after: N=7, n=16. (**F**) Dilation duration between the half-peaks. (**G**) The dilation amplitudes of vessel diameter change induced by WP stimulation, pinacidil, papaverine, PNU and control. WP stimulation: N=8; Pinacidil: N=9; Papaverine: N=8; PNU: N=4; Control: N=5. (**H**) Absolute diameter change upon WP stimulation, pinacidil, or papaverine. One-way ANOVA with post hoc test.

To examine whether K_ATP_ channels are involved in NVC, we compared WP stimulation-induced vasodilation before and after topical application for 45 minutes with PNU (**Fig. 4D – E**). This exposure significantly dampened the WP stimulation-induced vasodilation amplitude response of the PA, but not statistically significant of the capillaries although p values were close to 0.05 (**Fig. 4D – E**). PNU reduced the half-peak duration for all vessel categories, whereas the latency to half-peak increase remained unchanged for the PA (**Fig. 4G, Supplementary fig. 3B**). This result suggested that K_ATP_ channels may maintain and prolong WP stimulation-induced vasodilation via VSMC mechanisms in PA and 1–2 order capillaries.

To assess where pinacidil exerts its action in the dynamic spectrum of dilation, we compared the amplitude of the vasodilator responses to pinacidil with WP-induced vasodilation and the vascular effects of local puffing of papaverine. Papaverine relaxes VSMC by inhibiting breakdown of cGMP [25] and blocking voltage-gated calcium channels [26]. The response to papaverine is commonly used to document the preserved VSMC function [15]. Local papaverine and WP stimulation triggered dilation of the PA and 1–3 order capillaries that was comparable to the effect of pinacidil (**Fig. 4G**). This result suggested that at 5 mM pinacidil dilates the PA and capillaries within a physiological range.

NO may relax VSMCs or pericytes by activation of K^+^ channels in addition to the cGMP pathway. Although much of the evidence indicates that K-Ca channels are the primarily involved, NO activation of K_ATP_ channels also plays a role [27]. To probe this potential role in NO-associated pericyte relaxation, we topically applied the K_ATP_ channel blocker PNU before puffing SNAP. The *in vivo* vasodilation and *in vitro* pericyte calcium response induced by SNAP were the same with and without PNU, ruling out involvement of K_ATP_ channels in NO-induced pericyte relaxation *in vivo* (**Supplementary fig. 4**) and *in vitro* (**Supplementary fig. 5**).

We also examined whether pinacidil affected fEPSP and fIPSP amplitudes and latencies evoked by WP stimulation. For this purpose, we compared recordings taken 4 minutes before and 7 minutes after pinacidil puff (**Supplementary fig. 2G – I**), or before and after PNU perfusion (**Supplementary fig. 2J – L**). Both amplitudes and latencies of fEPSP and fIPSP remained constant. To directly rule out the possibility of K_ATP_ in neurons and astrocytes participating in pinacidil-induced vasodilation, we examined the neuronal and astrocytic calcium responses by local bulk loading of OGB Oregon Green BAPTA-1/AM (OGB) and sulforhodamine 101 (SR101) [28, 29]. Pinacidil puffing had no effect on neuronal or astrocytic calcium (**Supplementary material and supplementary fig. 6**). The results indicate that pinacidil does not affect neuronal or astrocytic [Ca^2+^]_i_ or fEPSPs and fIPSPs, and that it is unlikely that neurons or astrocytes indirectly mediate the pinacidil response, i.e., K_ATP_ channel–mediated vasodilation.

### K_ATP_ channels mediate vasodilation by Ca^2+^-dependent pericyte signaling

To characterize the mechanism by which K_ATP_ channels may affect pericytes, we used an *in vitro* preparation with pericyte monoculture from bovine brain. Immunohistochemical staining of the pericyte markers αSMA and platelet-derived growth factor receptor-β (PDGFR-β) were used to confirm the cell type (**Supplementary fig. 7A**). Pericytes were pre-loaded with 5 µM Fura-2 AM, a ratiometric indicator of intracellular Ca^2+^. Intracellular Ca^2+^ levels were measured in a plate reader during application of pinacidil at concentrations ranging from 0.67 µM to 67 µM. The pericytes responded to extracellular pinacidil by a significant decrease in cytosolic Ca^2+^ (**Supplementary fig. 7B, C**). This result is consistent with the assumption that opening of K_ATP_ channels triggers potassium outflow and hyperpolarization that blocks voltage-dependent L-type Ca2+ channels. To confirm this hypothesis, we preconditioned isolated pericytes with 67 µM PNU or 67 µM nifedipine (blocker of L-type Ca^2+^ channels) before addition of 20µM pinacidil. Both PNU and nifedipine dramatically attenuated the pinacidil-induced calcium drop in pericytes (**Supplementary fig. 7D, E**), confirming that pericyte L-type Ca^2+^ channels play a major role in the K_ATP_ channel-mediated decreases in pericyte Ca^2+^. Taken together, the data suggest that pericyte K_ATP_ channels are closed under physiological resting conditions but, open in response to an appropriate agonist. They appear to contribute to NVC under conditions of increased Na^+^-K^+^-ATPase activity, causing a reduction in the free concentration of intracellular calcium in contractile mural cells that results in local vasodilation.

### ET1-induced vasoconstriction is caused by contraction of pericyte somas and processes and mediated by inositol triphosphate receptor–dependent Ca^2+^ increases

ET1 is a vasoactive peptide released from the vascular endothelium that regulates blood pressure and vascular tone by promoting vasoconstriction. We examined pericyte and the vascular response to local puffing of ET1 on the PA and 1**–**3 order capillaries *in vivo* and *in vitro. In vivo* puffing of 0.5µM ET1 instantly triggered strong vasoconstriction (**Fig. 5A – B**). The constriction was long-lasting (> 9 min, **Fig. 5B, F**), and the most severely constricted vessel segments co-localized with pericyte somas (green and cyan lines) and adjacent regions devoid of soma (magenta line) at 1^st^ and 2^nd^ -order capillaries. Thus, our results suggest that both VSMCs on arterioles and pericytes on capillaries respond to ET1. Furthermore, preconditioning with the endothelin receptor type A antagonist BQ123 abolished ET1 induced vasoconstriction (**Fig. 5C–F**).

**Figure 5.**
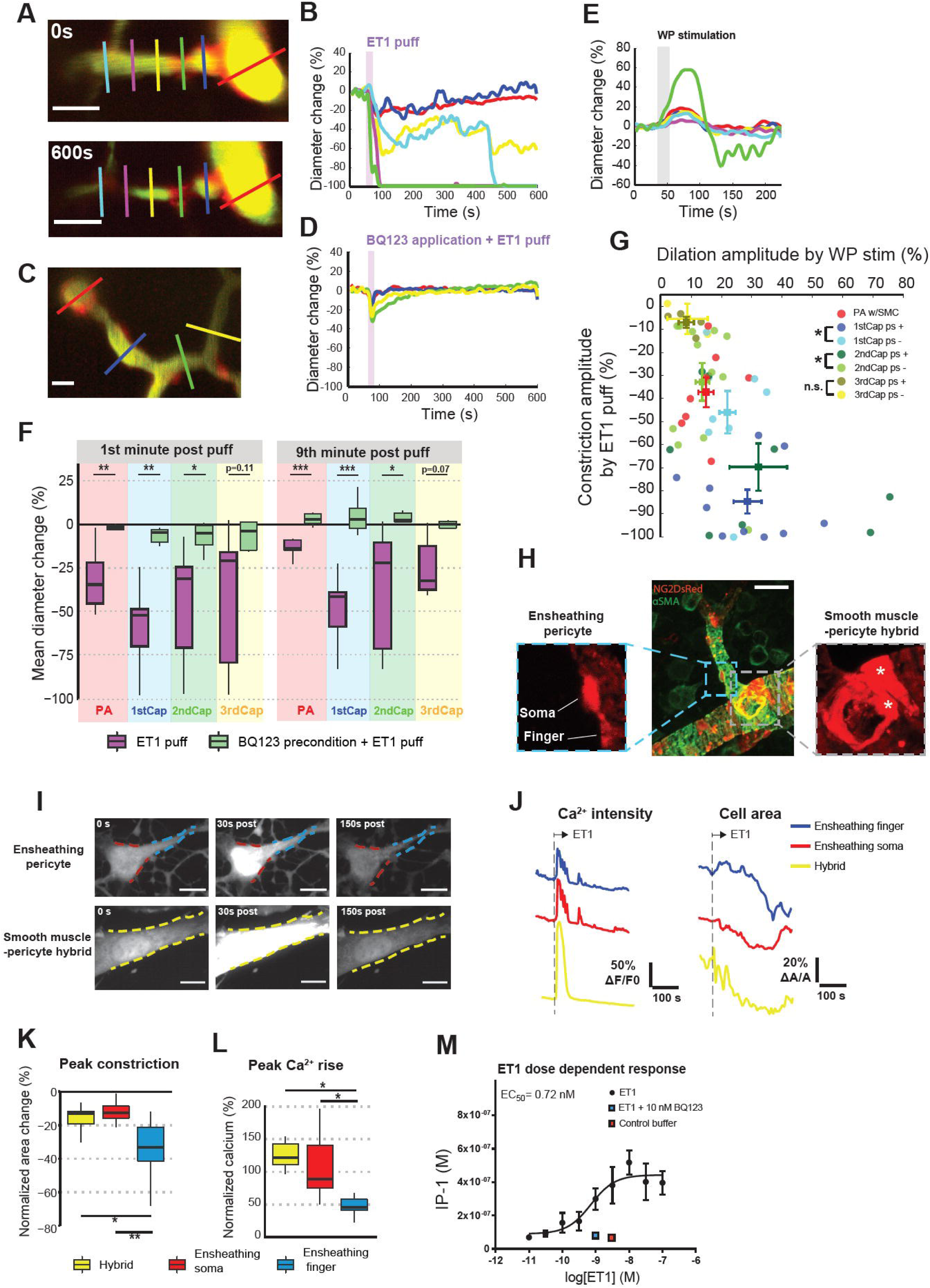
Endothelin-1 (ET1) induces contraction on both pericyte soma and processes. (**A**) Representative images of 0.0005 mM ET1-puff-induced vasoconstriction. Each image is averaged from a local image stack. Upper: resting state. Lower: 9 minutes after puff. Scale bar: 20 µm. (**B**) Normalized diameter changes over time at each color-coded ROI in (A). (**C**) Representative image of ET1 puff after topical application of 0.5mM BQ123 (ET1 receptor antagonist) for 45 minutes. (**D**) Normalized diameter changes over time at each color-coded ROI in (C). (**E**) Vasodilation curves induced by WP stimulation at the same ROIs with (A) before ET1 puff. (**F**) Comparison of mean diameter change at 1^st^ and 9^th^ minute post ET1 puff with or without BQ123. Without BQ123: N=5, n=6; with BQ123: N=6, n=11. (**G**) Co-registration of ET1-induced vasoconstriction and WP stimulation induced vasodilation at the same vessel segments (ROIs). The ROIs are classified as PA (red), 1^st^ (blue), 2^nd^ (green), 3^rd^ (yellow) order capillaries, and co-localization with pericyte soma (ps+) or without (ps-). Each dot represents an individual ROI. Error bars denote the mean ± SEM for the same color-coded ROI type. (**H**) Representative image of immunohistochemical staining αSMA in NG2DsRed mouse brain slices. Green: αSMA; red: pericytes. Based on morphology, contractile mural cells (αSMA positive and NG2 positive) are classified as ensheathing pericyte (left inset) or smooth muscle pericyte hybrid (right inset). Scale bar: 8 µm. (**I-J**) Representative responses of cultured pericyte upon ET1 administration. Based on time-lapse movies (I), normalized Ca^2+^ intensity and cell area changes are plotted in (J) from smooth muscle pericyte hybrid, ensheathing pericyte soma and finger process. Scale bar: 10 µm. (**K-L**) Statistical comparison of peak contraction (K) and Ca^2+^ rises (L) at hybrid, ensheathing soma and ensheathing finger. One-way ANOVA with post hoc test. (**M**) ET1 dose dependent IP1 responses. The precondition of BQ123 blocks the IP1 responses (blue square), comparable with the addition of the control buffer. Data show means ± SEM of 3–6 biological replicates.

Next, we asked whether vessel segments with the strongest vasoconstriction also responded most strongly to vasodilator stimuli and whether this pattern of high reactivity co-localized with pericytes. We classified the vessel segments (regions of interest; ROIs) as being with or without pericyte soma, but could not visualize pericyte processes at this stage because of the limitations of our two-photon microscope. For both dilation and constriction, 1^st^ and 2^nd^ order capillaries with pericyte somas showed stronger reactivity to ET1 than segments without pericytes (**Fig. 5G**), indicating an active role of pericyte in ET1-induced vasoconstriction. Finally, ET1 receptors did not contribute the WP stimulation–induced fEPSP and fIPSP response amplitudes (**Supplementary fig. 2M – O**), or dilation amplitude, duration and time to half-peak (**Supplementary fig. 8A – E**), suggesting an inactive role of ET1 in NVC.

### ET1-triggered vasoconstrictor signaling pathways in pericytes of various shapes

To examine the cellular basis of pericyte contraction, we used immunohistochemistry to stain αSMA in NG2DsRed cells in brain slices (**Fig. 5H, middle**, αSMA in green, NG2 in red). αSMA-positive mural cells exhibited different shapes on arterioles and on 1^st^-order capillaries. On arterioles, αSMA-positive mural cells were spindle-shaped with sparse, short processes, which is in accord with the so-called smooth muscle–pericyte hybrid in other studies [30, 31] (**Fig. 5H, right**). We used the term ‘hybrid’ for the mural cells on PA near the junction with the 1^st^-order capillary. αSMA-positive mural cells on the 1^st^-order capillary were star-shaped with thick finger-shaped processes and mesh-shaped thin processes that embraced the capillary wall (**Fig. 5H, left**), consistent with the term ‘ensheathing pericytes’ [31, 32], which we apply to them here. The thick processes of both ensheathing pericytes on capillaries and pericyte hybrids on arterioles expressed a high level of αSMA (green).

To assess the process of constriction, we examined changes in Ca^2+^ and pericyte shape at the soma and processes in cultured bovine pericytes loaded with the Ca^2+^ indicator Cal-520 AM. The cultured pericytes were also classified as ensheathing or hybrid based on their star or spindle shape, as detected by light microscopy. Immediately after administration of 50 nM ET1, calcium rose in pericytes regardless of shape, followed by contraction, both responses were reduced by preconditioning with BQ123 (**Fig. 5I-J, supplementary fig. 9**). Of note, thick processes of ensheathing pericytes contracted more strongly than the hybrids but displayed a smaller calcium increase (**Fig. 5K – L**). The data are consistent with those of our *in vivo* study, in which ET1-induced vasoconstriction was observed at some ROIs where pericyte somas were absent (**Fig. 5A**). Immunohistochemistry data confirmed co-expression of αSMA in soma and processes of ensheathing and hybrid NG2DsRed cells (**Supplementary fig. 10**). Calcium passing through Ca^2+^ channels opened by second messenger inositol triphosphate receptor (IP3) is the assumed mechanism of ET1-induced vasoconstriction, but whether it represents the mechanism in brain pericytes is unknown. IP3 is rapidly degraded to inositol biphosphate and then inositol monophosphate (IP1), the latter of which we used as an indicator for IP3-mediated calcium elevation. We stabilized IP1 using LiCl in cultured pericytes and measured an ET1 dose-dependent rise in IP1 (**Fig. 5M**) [33, 34]. These data suggested that IP3 mediates ET1-induced pericyte constriction of arterioles and capillaries and that both VSMC and pericytes respond to ET1 with contraction because of their high αSMA content.

### Endothelin receptors mediate pericyte contractions after ischemia

Global ischemia is accompanied by vascular constriction within minutes, and the conventional understanding of this reaction is related to the release of vasoconstrictors from the ischemic brain. It is also possible, however, that BECs release the vasoconstrictor, as they are loaded with mitochondria and assumed to be sensitive to lack of oxygen [35]. For this reason, we examined whether the contraction of pericytes in global ischemic stroke is associated with ET1 signaling [36]. VSMCs on PAs and pericytes at the 1^st^ and 2^nd^ order capillaries contracted instantly during the first 10 minutes after cardiac arrest (**Fig. 6A upper**), whereas pericytes at 3rd-order capillaries were only moderately constricted. Capillaries with visible pericyte somas exhibited stronger constriction than capillaries lacking pericyte somas (**Fig. 6A**). The vascular pattern of ischemia-induced vasoconstriction was like that induced by ET1. This finding led us to examine whether ET1 receptors were involved. For this purpose, we topically applied the cortex with the specific ET1-receptor antagonist BQ123 (0.5 mM) for 45 minutes before cardiac arrest. BQ123 mitigated the pericyte contraction in global ischemia (**Fig. 6A, lower**). The mitigation was co-localized with pericyte somas at the PA and 1^st^ and 2^nd^ order capillaries (**Fig. 6B**). Our result implies a role for ET1 in pericyte-associated capillary constriction in global ischemia [37].

**Figure 6:**
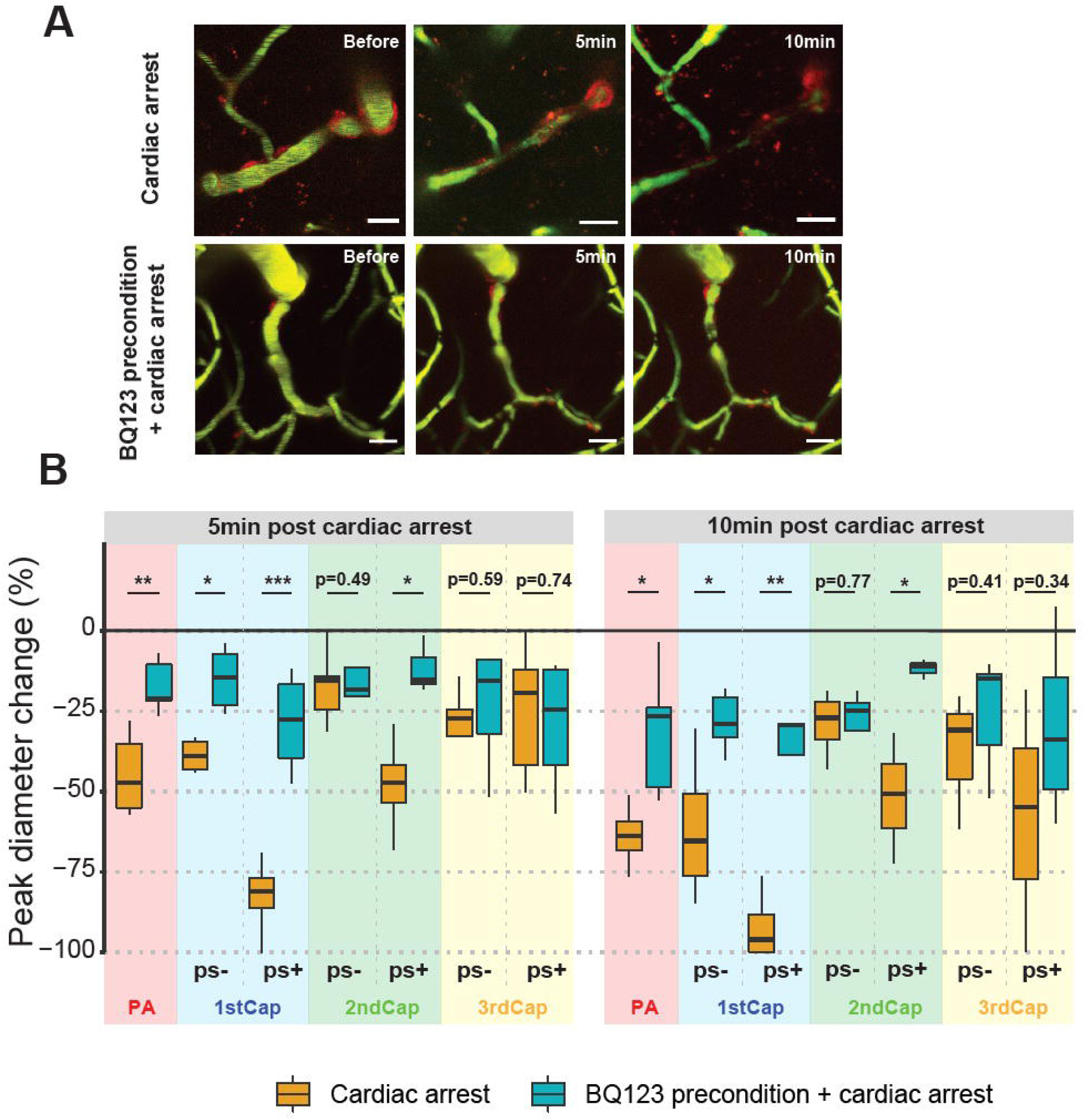
Pericyte contractions after ischemia are mediated by endothelin receptors. (**A**) Representative time-lapse images of PA and 1–3 order capillaries under cardiac arrest induced by i.v. bolus infusion of pentobarbital. Upper: normal cardiac arrest; lower: precondition of BQ123, which mitigated ischemia-induced pericyte constriction. Scale bar: 20 µm. (**B**) Summary of diameter change at PA and 1–3 order capillaries co-localized with pericyte somas or without, upon cardiac arrest with or without BQ123 precondition. ps- and ps+ denote without and with pericyte some respectively. Without BQ123: N=6, n=9; with BQ123: N=5, n=10.

### 1^st^ order capillaries are the main gatekeepers of capillary blood flow

Throughout our experiments, we noted that strongest capillary responses occurred at the 1st order capillary This result is in accordance with our previous observations of conducted vascular responses and of a precapillary sphincter in the vascular segment between the PA and the branch point of the second-order capillary. To assess the possible implications of a special role of the 1^st^ order capillary, we developed a mathematical model based on construction of a cerebrovascular network encompassing a PA, capillaries up to the third branching order, and a penetrating venule. The model assumed that flow resistances were consistent with Poiseuille’s law and that the inlet pressure of the PA was 25 mmHg and the venous outlet pressure was 5 mmHg. This setup allowed us to calculate network blood flow and pressure using Kirchoff’s 2^nd^ law (**Fig. 7A**; insert depicts first proximal branching of the PA and associated downstream capillaries). From the resting state (control), we assessed how blood flow changes correlated with relative diameter changes in individual capillaries, with all else being equal.

**Figure 7:**
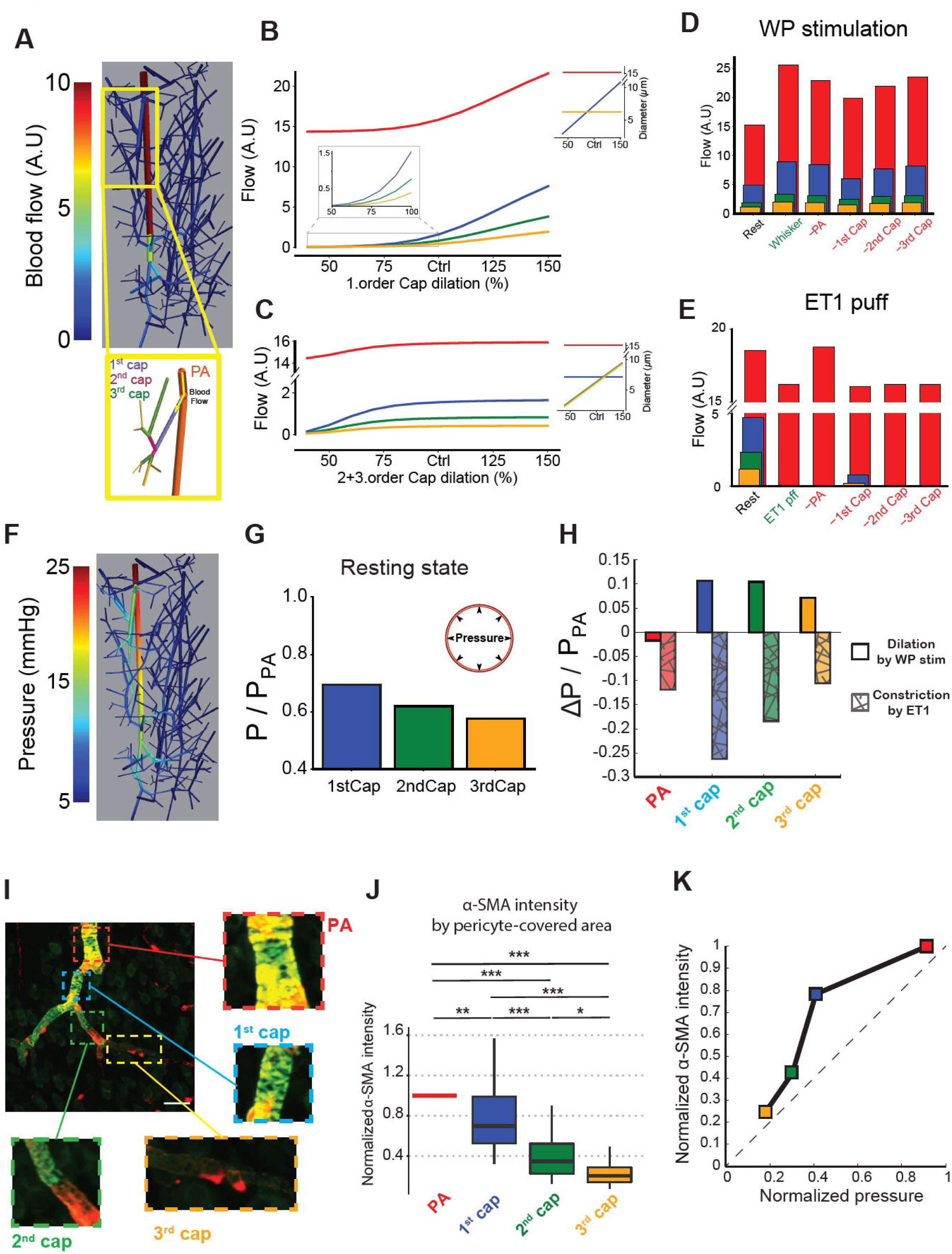
1^st^ order capillaries are main gatekeepers of capillary blood flow. (**A**) Mathematical modeling of flow distribution in a reconstructed cerebrovascular tree containing one penetrating arteriole and venule, respectively (see Methods), under resting conditions. (**B–C**) Summary of the calculated blood flows in the vessels (PA: red; 1^st^ : blue; 2^nd^ : green; and 3^rd^ order capillary: orange) under resting conditions as either the diameter of the 1^st^ order capillary (B) or diameters of the 2^nd^ and 3^rd^ order capillaries (C) (see inserts) were changed relative to control (Ctrl). (**D**) Summary of calculated average blood flows among all PA, 1–3 order capillary segments in the network before (rest) and during WP stimulation. The groups labeled in red denote calculations of WP stimulations in which either the PA, 1–3 order capillary segments were disallowed from dilating. Note the 1^st^ order capillary has the strongest influence on network flows. (**E**) Summary of calculated blood flows in the vessels around the 1^st^ proximal branch of the PA before (Rest) and during ET1 puff. As in (E), calculations of the puff where segments were disallowed from constricting are shown in the red groups. (**F**) Mathematical modeling of pressure distribution in the resting state. Inlet pressure into the PA was set to 25 mmHg, whereas all outlet pressures were set to 5 mmHg. (**G**) Summary of the average pressures in all 1^st^, 2^nd^, and 3^rd^ order capillaries in the network normalized to the average pressure in the PA. (**H**) Normalized pressure change at PA, 1–3 order capillaries by dilation or constriction. (**I**) Representative image of αSMA staining among the PA (red), 1^st^ order capillaries (blue), 2^nd^ order capillaries (green), and 3^rd^ order capillaries (orange), see inserts for enlargements. (**J**) Summarized intensity of αSMA in pericytes on 1^st^, 2^nd^, or 3^rd^ order capillaries normalized to the intensity from smooth muscle on the PA. Note that αSMA is significantly more expressed in 1^st^ order capillaries than 2^nd^ and 3^rd^ order; whereas all 1– 3 order capillaries contained less αSMA than PA. N=3, n=31. (**K**) Relationship between normalized pressure and normalized αSMA density at peak dilation upon WP stimulation.

An increase of the diameter of all 1^st^ order capillaries (**Fig. 7B**) in the network strongly increased the blood flow through the PA, 1^st^, 2^nd^, and 3^rd^ order capillaries (red, blue, green, and orange curves, respectively). In contrast, simultaneous dilation of both 2^nd^ and 3^rd^ order capillaries without dilation of the 1^st^ order capillary did not augment blood flow (**Fig. 7C**). As expected, constriction of either 1^st^ order capillaries or the 2^nd^ and 3^rd^ order capillaries strongly reduced blood flow in the capillaries. The results suggested that the 1^st^ order capillary gates blood flow under resting conditions [9, 10, 38].

Next, we calculated the average blood flow for WP stimulation (**Fig. 7D**, green group) using the observed diameter changes in vessel segments (**Fig. 4H**). We compared this flow pattern to rest (black group) and to hypothetical scenarios where the diameters of either the PA, 1^st^, 2^nd^, or 3^rd^ order capillaries were kept at rest, i.e., could not dilate (red groups). Compared to rest, blood flow through the 1^st^ order capillary almost doubled during WP stimulation. Disabling vessel dilation of each segment decreased capillary blood flow during WP stimulation. Yet, the largest decrease (32.6%) was observed when the 1^st^ order capillaries were kept at the resting diameter (compare blue bars in **Fig. 7D**). A similar set of blood flow calculations during local ET1 constriction (**Fig. 5F**) of the network around the first proximal branch of the PA is shown in **Fig.7E.** ET1 puffing effectively reduced local capillary blood flow. However, capillary blood flow was partially restored by disabling constriction at the 1^st^ order capillary. Collectively, the results of these simulations support the hypothesis that capillary blood flow depends mainly on the diameter of 1^st^ order capillaries and that downstream capillaries contribute less to capillary flow control.

We next compared the pressure distribution in the network model under resting conditions (**Fig. 7F**). The average pressures in the capillaries at rest (summarized in **Fig. 7G**) suggest that the 1^st^ order capillary must withstand more pressure than downstream capillaries but less than the PA. Both during WP stimulation and during ET1 puff, the 1st order capillary is exposed to the largest pressure changes (**Fig. 7H**). The ability of 1^st^ order capillaries to produce the strongest capillary flow changes and maintain contractile ability in the face of large pressure changes, is expected to be matched a large contractile capacity of capillaries closest to the PA. We stained for αSMA, a central contractile component in vascular mural cells, to assess the relative expression in cortical capillaries and identified the largest expression among pericytes at the 1^st^ order capillary (**Fig. 7I-J, supplementary fig. 11**). This finding is consistent with both the calculated pressure distribution and the observed flow changes of different capillary segments (**Fig. 7B-E**). Very low level of αSMA was observed in 3^rd^ or higher order capillaries consistent with these capillaries having very limited control over capillary blood flow.

Finally, we plotted the normalized pressure during WP stimulation against the normalized αSMA expression as a proxy for force generating ability (**Fig. 7K**). The 1^st^ order capillary data point is furthest away from the identity line, indicating that the 1^st^ order capillary has the highest contractile capacity of all capillaries and that this vascular segment is the main regulator of capillary blood flow under physiological conditions.

## DISCUSSION

We measured neural activity and vascular dynamics by *in vivo* two-photon microscopy to identify mechanisms underlying endothelial-pericyte crosstalk and NVC in the mouse brain *in vivo*. We discovered that (1) capillary endothelial cells via the eNOS and ET1 pathways contribute to regulation of VSMCs and central nervous system pericytes and thus to arteriolar and capillary vascular tone; (2) that NO acts as a vasodilator via cGMP release that produces pericyte relaxation; (3) that activation of K_ATP_ channels in pericytes produce vasodilation mainly of 1^st^ -order capillaries and enhance and prolong the NVC response; and (4) that the mechanism underlying this effect is associated with closing of L-type Ca^2+^ channels at the pericyte membrane. Furthermore, we show that (5) ET1-induced pericyte contraction occurs at the soma and at thick, finger-shaped processes, via IP3 signaling and (6) that capillary blood flow is regulated primarily by pericytes at 1^st^ order capillaries as a result of higher expression of αSMA in 1^st^ order capillaries than in pericytes at downstream capillaries.

K_ATP_ channels exist in almost all cell types in the brain, including endothelial cells, VSMCs, neurons and astrocytes. Recent findings have revealed that K_ATP_ channel expression is several magnitudes higher in brain pericytes than in other cell types [39, 40]. Our findings suggest that K_ATP_ channels in pericytes enhance and prolong the vasodilator response to increased brain activity. Similar to VSMCs [41], the signaling pathway involves potassium outflow and membrane hyperpolarization, which inactivates L-type Ca^2+^ channels and leads to vasodilation through reduced intracellular Ca^2+^.

In big arteries and arterioles, the NO/cGMP signaling cascade mediates endothelium-dependent relaxation of smooth muscle cells [42]. Here, we demonstrated that the NO/cGMP pathway is also a signaling pathway from endothelium to pericytes in brain capillaries; however, we did not confirm a primary involvement of the pathway in NVC. These results are consistent with those of studies in rats [43, 44] and some in mice [4], but other mouse studies have shown that NVC can be curtailed by L-NAME infusion in wild-type mice and in eNOS knockout mice [45]. The discrepancy could trace to different species or strains (rats versus mice, different mouse strains) or locations (capillary in our study versus previous reports on artery/arterioles). That said, it is important to appreciate that L-NAME does not cross the blood–brain barrier and that the NO-cGMP pathway modulates rather than mediates NVC in the somatosensory cortex [46]. Therefore, the absence of an L-NAME effect may be consistent with a greater relevance of other modulators. In comparison, blocking cGMP synthesis with ODQ led to a strongly decreased dilation amplitude and duration by WP stimulation, which is consistent with neurotransmitter-mediated release of signaling molecules, NO or CO, which activates sGC in vascular mural cells [47].

We examined the cellular signaling mechanisms of pericyte contractility and capillary diameter regulation with a focus on ET1 and NO and the K_ATP_ signaling pathways. In VSMCs and pericytes, increases in free intracellular calcium by ET1 activates myosin light chain kinase, which facilitates αSMA contraction and thus vasoconstriction [48, 49]. In comparison, NO activates the sGC and increases cGMP, resulting in a calcium decrease, which activates myosin light chain phosphatase, facilitating vasodilation [50]. cGMP-dependent protein kinase G (PKG), which lowers free intracellular calcium by inhibition of IP3 receptors, activation of the sarco/endoplasmic reticulum Ca^2+^-ATPase (SERCA) pump and calcium clearing out of the cell, is also activated by cGMP [51].

In the last part of our study, we asked why the 1^st^ order capillaries exhibit most profound vasoresponses? They are perfectly located to control local capillary blood flow and supply energy to active nerve cells. 1^st^ order capillaries generate the largest response to both systemic infusion of L-NAME, WP stimulation and local stimulation (puff of pinacidil, Ach, SNAP and ET1). This led us to hypothesize that pericytes at 1^st^ order capillaries have larger contractility than at 2^nd^ order capillaries. We show that αSMA at pericytes of 1^st^ order capillaries was less abundant than for VSMC and pericyte hybrids at the PA, but more abundant than for pericytes at 2^nd^and 3^rd^ order capillaries. This pattern of αSMA expression fits with the modeled pressure profile among vessel segments. The arterial pressure head in the systole is strong at the PA, which must be able to produce a great force in order to maintain a contractile ability. Indeed, our model of the pressure distribution in a reconstructed vascular network showed a pattern of pressure distribution that matched αSMA expression (**Fig. 7**). The model also assessed how blood flow may be distributed among the PA, 1^st^ order, 2^nd^ order, and 3^rd^ order capillaries with diameter changes of each vessel segment. During rest, capillary blood flow will increase only in response to dilation of the 1^st^ order capillary. Diameter changes in 2^nd^ or higher order capillaries are mainly passive and of little consequence for flow in the capillary network. Our data are in agreement with previous studies that detected αSMA in pericytes up to the 3^rd^ – 4 ^th^ order of capillaries, and with the notion that pericyte regulation of capillaries is mechanistically related to αSMA contraction and relaxation [32, 52].

In summary, our data provide mechanistic insight into the integrated vascular function that regulates brain capillary blood flow by identifying distinct cellular mechanisms evoked by three key signaling pathways in brain pericytes in the living brain. These results reveal a strong interaction between BECs and pericytes and a special regulatory role for pericyte K_ATP_ channels. In cardiac arrest, the endothelial vasoconstrictor ET1 may be a key signaling molecule causing persistent capillary vasoconstriction and resulting in impaired microvascular reperfusion. Despite the slow response time of pericytes, their response amplitudes can reflect the underlying neuronal activities and could reveal interactions among different vascular mechanisms. The results of this work have the potential to increase understanding of the complexity of vascular regulation as a result of the interplay between signals from endothelial cells and nerve cells. In addition, these findings have translational implications for a better understanding of cerebrovascular and neurodegenerative disorders. The results expand the boundaries in the universal effort to comprehend how integrated vascular networks may work to match the brain’s energy demand.

## ACKNOWLEDGMENTS

We would like to acknowledge Assistant Professor Krzysztof Kucharz and Associate Professor Henrik Flyvbjerg for scientific discussions. We acknowledge Assistant Professor Søren Grubb for providing z-stack for our modeling work. We acknowledge the Core Facility for Integrated Microscopy, Faculty of Health and Medical Sciences, University of Copenhagen, were we used confocal and spinning disc confocal microscopy in our *in vitro* studies. We further acknowledge Dr. Thor Møller for introduction to- and assistance with the IP-one assay. This study was supported by the Lundbeck Foundation, the Danish Medical Research Council, the Alice Brenaa Foundation, Augustinus Foundation, Carl og Ellen Hertz Familielegat and Nordea Foundation Grant to the Center for Healthy Aging.

## AUTHOR CONTRIBUTIONS

M.J.L. and B.B. designed research; S.A.Z., H.C.C.H., J.C.F., C.C. and M.L. performed research; C.C. contributed analytic tools; S.A.Z., C.C., H.C.C.H., B.O.H. and J.C.F. analyzed data; B.O.H. provided math models; C.C., S.A.Z. and M.J.L. wrote the paper.

The authors declare no conflict of interest.

## DATA AVAILABILITY

The data that supports the findings of this study are available from the corresponding author upon request.

## METHODS

### Animal handling

All procedures were approved by the Danish National Ethics Committee according to the guidelines set forth in the European Council’s Convention for the Protection of Vertebrate Animals used for Experimental and Other Scientific Purposes and are in compliance with the ARRIVE guidelines. A total of 11 C57bl6/J and 65 NG2DsRed mice Tg (cspg4-DsRed.T1)(1Akik/j) from the Jackson laboratory), age 3–12 months and of both sexes, were used. We cannulated the mouse trachea for mechanical ventilation. One catheter was placed into the left femoral artery to monitor blood pressure and blood gases; a second catheter was inserted into left femoral vein for infusion of substances. To ensure that the animals were kept under physiological conditions, we continuously monitored end-expiratory CO_2_, blood pressure, heart rate and O_2_ saturation at the right hind paw. Furthermore, we assessed blood gases in arterial blood samples twice during an experiment (pO_2_, 95–110 mmHg; pCO_2_, 35–40 mmHg; pH, 7.35–7.45). Body temperature was maintained at 37°C using a rectal temperature probe and heating blanket. A craniotomy was drilled with a diameter of ∼4 mm, centered at 0.5 mm behind and 3 mm to the right of bregma above the sensory barrel cortex region. After dura removal, the preparation was covered with 0.75% agarose gel, moistened with artificial CSF (aCSF; NaCl 120 mM, KCl 2.8 mM, NaHCO_3_ 22 mM, CaCl_2_ 1.45 mM, Na_2_HPO_4_ 1 mM, MgCl_2_ 0.876 mM, and glucose 2.55 mM; pH=7.4), and kept at 37°C. For imaging experiments, part of the craniotomy was covered with a tilted glass coverslip that permitted the insertion of micro-electrodes and pharmacological interventions. A detailed description has been published in [14]. The mice were anesthetized by intraperitoneal injection of xylazine (10 mg/kg) followed by ketamine (60 mg/kg), and maintained during surgery with supplemental doses (30 mg/kg) of ketamine every 24 minutes. Upon completion of all surgical procedures, anesthesia was switched to continuous i.v. infusion with a mixture of 17% α-chloralose and 2% fluorescein isothiocyanate (FITC)-dextran (0.02 mL/10 g/h). At the end of the experimental protocol, the mice were euthanized by i.v. injection of 0.05 mL pentobarbital followed by cervical dislocation.

### Whisker-pad stimulation

The mouse sensory barrel cortex was activated by stimulation of the contralateral ramus infraorbitalis of the trigeminal nerve using a set of custom-made bipolar electrodes inserted percutaneously. The cathode was positioned corresponding to the hiatus infraorbitalis (IO), and the anode was inserted into the masticatory muscles [53]. WP stimulation (thalamocortical IO stimulation) was performed at an intensity of 1.5 mA with pulse duration of 1 ms, in trains of 20 s at 2Hz.

### Local ejection of substances (puffing) by glass micropipette

Borosilicate glass micropipettes were produced using a pipette puller (P-97, Sutter Instrument), with a resistance of 3–3.5 MΩ. The pipette was loaded with a mixture of 10 µM Alexa 594 and vasoactive substances, which enabled visualization of pipette tip under an epi-fluorescent camera and two-photon microscope. Under both epi-fluorescent imaging and two-photon microscopy guidance, the pipette was carefully inserted into the cortex and approached the proximity of the targeted vascular region. The distance between the pipette tip and the vasculature was 30–50 µm (**Fig. 1B**). Pressure ejection of vasoactive substances was performed using an air pressure pump at 8–15 psi. A red cloud (Alexa 594) ejected from the pipette tip was visually observed to cover the local vascular region instantaneously and simultaneously, and the background returned to normal approximately 1 minute after puffing [9]. An electrode inside the puffing pipette was used to record local electrical brain activity during the experiments.

### Bulk-loading of OGB and SR101

Ca^2+^ **-**sensitive dye Oregon Green BAPTA-1/AM (OGB; Invitrogen, solubilized in 20% pluronic acid and dimethylsulfoxide and diluted to 0.8 mM with aCSF) and the astrocyte-specific marker SR101 (Sigma-Aldrich, solubilized in methanol and diluted to 0.05 mM in aCSF) were co-injected by glass micropipette into the mouse whisker barrel cortex at a depth of 100–200 μm (**Supplementary fig. 6A**). The glass micropipette (impedance 1–1.5 MΩ) was connected to an air pressure pump with pressure of 15–20 psi and ejecting duration of 5–10 s.

### Two-photon imaging

FITC-dextran (MW 500,000, 50 µL, Sigma-Aldric) was applied i.v. to stain the vessel lumen as green color. Before two-photon imaging, 4% w/v FITC-dextran was administered as a bolus into the femoral vein to label the blood plasma. During the two-photon imaging, 2% of FITC-dextran was continuously infused at a slow speed to compensate for the metabolic loss of FITC-dextran (see also the animal handling session). Fast repetitive hyperstack imaging (4D imaging) was performed using a commercial two-photon microscope (Femto3D-RC, Femtonics Ltd.) and a 25 × 1.0 NA piezo motor objective. This method compensates for focus drift and allows for evaluation of vasculatures spanning a certain z-axis range. Each image stack was acquired within 1 second and comprised 10–14 planes with a plane distance of 3–5 µm. This approach covered the whole z-axis range of the investigated blood vessels. The pixel sizes in the x-y plane were 0.2– 0.38 µm. The excitation wavelength was set to 900 nm. The emitted light was filtered to collect red and green light from DsRed (pericytes) and FITC-dextran (vessel lumens), respectively.

### Data analysis

The imaging analytical tool was custom-made using MATLAB. For measuring vessel diameter in hyperstack video, each image stack was flattened onto one image by maximal intensity projection, and thus create a time-lapse movie. An averaged image over time from the green channel was plotted for ROI placement. Rectangular ROIs with a width of 4 µm was drawn perpendicularly across the vessel longitude (**Fig. 2A**). The rectangular ROI was averaged by projection into one line for each frame, representing the profile of the vessel segment at this frame. The profile line was plotted as a 2D image with the x axis as number of frames. Chan-Vese segmentation or pixel-intensity based segmentation was used to delineate the vessel edge and estimate the diameter change in the vessel segment at this ROI [9, 54, 55]. Less than 2% of the evoked diameter change was considered as non-responding but included in analysis as zero values. Up to 5 ROIs with adjacent distance of 10 µm could be placed at the same order capillary, depending on the length of this capillary order. Diameter curves at the same order capillary are averaged, representing the response curve at this capillary order. The vessel response amplitude was defined as the largest vasodilation/constriction amplitude after WP stimulation/puffing. The response latency was defined as the latency of half-max amplitude. The response duration was defined as the period between half-peaks of uprising and falling phases.

For measuring the calcium signals in neurons and astrocytes loaded by OGB and SR101, each image stack was flattened onto one image by average intensity projection. The purpose of using average intensity projection is to iron out the false calcium surge by a single super bright pixel. For each converted time-lapse movie, free-drawn ROIs were placed around neurons and astrocytic processes and end-feet. The labeling pattern of the SR101 dye was used to select ROIs as either astrocytic or neuronal. For each frame, pixels with SR101 intensities >1.5 standard deviations of the mean intensity in ROIs were selected as astrocytic, and the OGB intensity of these pixels averaged to create a time trace (F(t)) for every ROI. Evoked Ca^2+^ responses were detected if the intensities increased by ≥5% and ≥2 SD from baseline, with a duration ≥7 s, during a 60-second period starting from stimulation onset. Responsivity was defined by the fraction of ROIs responding to stimulation and reported as a ratio (%). For each ROI, mean intensity increases during the response were used as a measure of response size, reported as ΔF/F_0_ (%).

The evoked LFPs were recorded and measured using the software Spike2 v7.02a (CED, Cambridge Electronic Design). LFPs were overlaid and averaged by aligning stimulation marks. The evoked negative field potential was considered to be the fEPSP and the following positive potential as the fIPSP (**Supplementary fig. 2**). The fEPSP and fIPSP amplitudes are defined as the peak amplitudes from baseline and the latencies as from stimulation mark onset to the peak of the fEPSP and fIPSP.

### Drug application

We locally ejected vasoactive substances (pinacidil, PNU, papaverine, Ach, SNAP, endothelin) via glass micropipette. Pinacidil monohydrate (P154, Sigma Aldrich) and PNU 37883 hydrochloride (Cat. No. 2095, Tocris) were solubilized in dimethylsulfoxide and further dissolved with aCSF. Ach (Cat. No. 2809, Tocris), ODQ-CAS 41443-28-1 -Calbiochem (495320, Sigma Aldrich), ET1 (Cat. No. 1160, Tocris) and BQ-123 (Cat. No. 1188, Tocris) were directly dissolved in aCSF. Aliquots were stored at -20°C. L-NAME (N5751, Sigma Aldrich) was dissolved in saline and stored at -20°C. L-NAME at a dosage of 30 mg/kg was injected as a bolus i.v. SNAP (Cat. No. 0598, Tocris) was freshly dissolved in aCSF during each experiment. PNU, ODQ and BQ123 were applied topically on the cortex for 45 minutes before experiments were continued. The control puff solution consisted of aCSF and dimethylsulfoxide and was stored at -20°C. Before each experiment, each drug solution used for puffing was mixed 50/50 with 20 mM Alexa 594 (dissolved in aCSF). Concentrations in final puffing solution were as follows: pinacidil 5 mM, PNU 2.5 mM, Ach 2 mM, SNAP 5 mM, papaverine 10 mM, and ET1 0.0005 mM. The concentrations of solutions for topical application were PNU 0.5 mM, ODQ 0.5 mM and BQ-123 0.5 mM.

### Immunohistochemistry

Adult NG2-dsRed mice were transcardially perfused with 4% paraformaldehyde and their brains extracted and stored in paraformaldehyde for 4 hours. The tissues were then cryoprotected in phosphate-buffered saline (PBS) with 30% sucrose and 0.1% sodium azide for 48 hours, rapidly frozen on dry ice–chilled isopentane, and sectioned at 50 μm thickness using a cryostat. Sections were rinsed for 5 minutes three times in 0.1 M PBS. The sections were permeabilized and blocked in 0.5% Triton-X 100 in 1× PBS (pH 7.2) and 1% bovine serum albumin overnight at 4°C. Sections were further incubated for two nights at 4°C with mouse ACTA2-FITC antibodies (1:200; Sigma;F3777) in blocking buffer containing 1 to 5% bovine serum albumin in 0.25–0.5% Triton-X 100 in 1× PBS. The sections were then washed for 5 minutes three times in 0.1 M PBS and incubated in Hoechst (1:6000) for 7 minutes, rinsed again (3×5 min) one time in blocking buffer and two times in 1× PBS, and mounted using SlowFade™ Diamond Antifade Mountant (Invitrogen; S36963). Fluorescence images were acquired with a confocal laser scanning microscope (LSM 700) equipped with Zen software and 20×/0.8 NA and 63×/1.40 NA oil DIC M27 objectives at 1× (0.170 μm/pixel) and 4× (0.021 μm/pixel) digital zoom, respectively. Care was taken to ensure similar fluorescence across images.

### Quantification of αSMA in immunohistochemical brain slices

The NG2DsRed mouse brain slices stained with αSMA displayed different pericyte phenotypes and αSMA density in different order capillaries (**Supplementary Fig. 11A**; pericytes, red; αSMA, green). The existing problem of using immunohistochemistry to quantify protein density is the uneven fluorescent intensity of antibodies at different depths of brain slices, as antibody binding efficacy decreases towards the center of the slice. The second reason is confocal laser power reduces with imaging depth. To overcome this problem, we implemented a new image processing algorithm. The same background area of the high-resolution image stack was manually selected (**Supplementary Fig. 11B, left**). We then calculated the mean background intensity of each image plane and normalized the whole image plane to the peak value of background within the stack (**Supplementary Fig. 11B, middle**). The new image stack was then flattened by maximal intensity projection **(Supplementary Fig. 11B, right**). We used two methods to quantify the αSMA intensity: (1) by the whole vessel area in the αSMA image, normalized to the intensity of PA (**Supplementary Fig. 11C-D**); and (2) by the selected region in the αSMA image–based pericyte-covered area in the DsRed image and normalized to the intensity of PA (**Supplementary Fig. 11E-F**).

### *In vitro* pericyte study

#### Isolation and culture of bovine brain capillaries to obtain bovine brain pericytes

Bovine brains were acquired at the local abattoir (Mogens Nielsen Kreaturslagteri A/S, Herlufmagle, Denmark), and capillaries were isolated as reported in detail elsewhere [56]. To obtain pericyte-enriched cultures, the capillaries were seeded in uncoated culture flasks and cultured until confluent in Dulbecco’s Modified Eagle’s Medium (DMEM-AQmedium, Sigma-D0819) supplemented with 10% fetal bovine serum, 1% (v/v) MEM non-essential amino acids (×100), and 100 U/mL:100 μg/mL penicillin:streptomycin solution. When confluent, the cells were passaged twice, with an initial brief trypzination to remove loosely adhering cells followed by a stronger trypzination to harvest pericytes. The pericytes were washed in culture medium and frozen in liquid nitrogen in FBS+DMSO (9:1).

#### Calcium measurements in pericytes

Pericytes were seeded at a density of 31,000 cells per cm^2^ in a black 96-well plate (Corning 3603) and cultured for 4 days. The pericytes were loaded with a 5 µM Fura-2 AM calcium indicator, and intracellular calcium quantification was performed as previously described [9]. Antagonists (PNU 37997, nifedipine, and ODQ) were added prior to pinacidil or SNAP to ensure inhibition. For live cell imaging, pericytes were cultured on glass coverslips (#1.5, Gerhard Menzel B.V. & Co, Braunschweig, Germany) at a density of 21,000 cells per cm^2^. The cells were loaded with 14 μM Cal-520 AM (AAT Bioquest, Sunnyvale, CA, USA) following the same procedure as with Fura-2. The cells were examined by spinning disc microscope (Zeiss Cell Observer) to measure the cell area and calcium change upon ET-1 administration. Concentrations of 10 nM and 50 nM ET-1 assay buffer were perfused with or without BQ123 precondition for 20 minutes. Time-series movies were recorded at 1 Hz using 20× and 63× objectives. Each condition was tested in 4 wells within each experiment and the experiments were run as 6 biological replicates.

#### Measurements of IP1 generation

IP1 was detected using homogeneous time resolved fluorescence technology, as previously described [57]. Cells were seeded and cultured as for the calcium measurements. After 4 days the cells were incubated one hour with ligands, subsequently lysed in cell lysis buffer from the IP-One-Gq kit (Cisbio Bioassays, Codolet, France) and frozen to -20°C. Cell lysates were transferred to a 384 OptiPlate (Perkin Elmer, Waltham, MA, USA) and mixed 1:1 with antibody mixture from the IP-ONE-Gq kit, IP1-d2 and Anti-IP1-Cryptate diluted 1:40 in HBSS + 20 mH HEPES. The plate was incubated protected from light for 60 minutes at room temperature, and the fluorescent emission was read at 665 nm and 620 nm in an EnVision^®^ Xcite Multilabel Plate Reader (PerkinElmer, Waltham, MA, USA). IP1 content was calculated based on a standard curve covering concentrations from 2.7–11,000 nM according to the IP-ONE kit protocol.

#### Data analysis

The fluorescent ratios at excitation 340/380 nm were standardized against the average ratio in Fura-2–loaded cells with the addition of blank assay buffer. Fluorescence changes were normalized against the baseline prior to agonist addition. The time-series movies were analyzed for pericyte calcium and area changes in a custom-made MATLAB program. Log dose-response for IP1 generation was determined using Graphpad Prism version 7 (San Diego, CA, USA) and EC50 was calculated based on an agonist curve with variable slope. One-way ANOVA was applied to evaluate significant differences (α=0.05) followed by either Dunnett’s multiple comparisons test (α=0.05) to identify groups significantly different from the control group (blank assay buffer) or Tukey’s multiple comparison test (α=0.05) to evaluate significant differences among all tested groups. All statistics concerning the Fura2 and IP-1 assays were performed in Graphpad Prism version 7.

#### Computational modeling

We developed a simple hemodynamic network model based on image reconstruction of a single PA and venule and their associated capillaries. Briefly, Amira software (version 6.1) was used to segment a z-stack of images, preprocessed with a Gaussian filter and background subtraction to facilitate segmentation. Network nodes, edges, and vessel lengths and radii were extracted from the reconstruction. However, vessel radii of the upper PA and the 1^st^, 2^nd^, and 3^rd^ order capillaries were taken from original image data and for each simulation, the vessel radii of specific vessels in the network were deliberately changed. Assuming that the vessels are rigid and the flow (*Q*) is laminar, the flow resistances of individual vessel segments were calculated using Poiseuille’s law and the standard hemodynamics measure of flow resistance (*R*):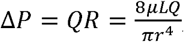, i.e.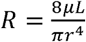, where P is pressure, *µ* is dynamic viscosity, *L* is vessel length, and *r* is vessel radius. In a network, Kirchoff’s law states that the sum of flows entering and leaving any internal node equals zero, 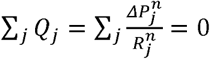 where 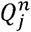 is the flow, 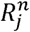 is the vascular flow resistance and 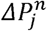 is the pressure drop in the *j*’th vessel entering the *n*’th node. We applied the empirical model describing the changes in apparent viscosity of blood (μ) with diameter (D) and hematocrit [58, 59]:

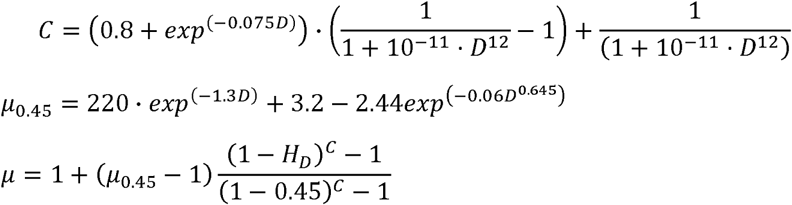

The discharge hematocrit (H_D_) was calculated based on a tube hematocrit (H_T_) of 0.3 [60, 61]:

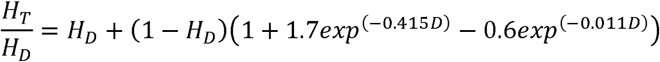

To solve the system of linear equations, we chose the boundary conditions such that inlet pressure into the PA was 25 mmHg in the control situation and every outlet pressures of the capillaries was 5 mmHg. The system was solved using the root solver in SciPy (1.1.0).

#### Limitations

All assumptions underlying Poiseuille’s law and Ohmic resistances in laminar flows apply. The boundary conditions strongly influence the solution because the system is forced to comply with the preset boundary pressures. However, the effects of changing diameters on the pressure and flow within the network are evident. We also assume that the empirical formulas to calculate blood viscosity apply to the cerebral microcirculation of mice.

### Statistical analysis of *in vivo* studies

Datasets are presented as boxes with whiskers. The box displays a 95% confidence interval around the median based on assumption of normal distribution. The median is marked as a thick horizontal line. The upper and lower whiskers extend to the largest and smallest values, respectively, within the 1.5× interquartile range. P values are from one-way ANOVA with Tukey-Kramer post hoc test or unpaired or paired Student’s t-tests, as appropriate. * indicates p<0.05, ** indicates p<0.001, *** indicates p<0.0001, and n.s. indicates not significant. All statistical analyses were performed using R (version 3.4.4).

## FIGURE LEGENDS

**Supplementary figure 1.**
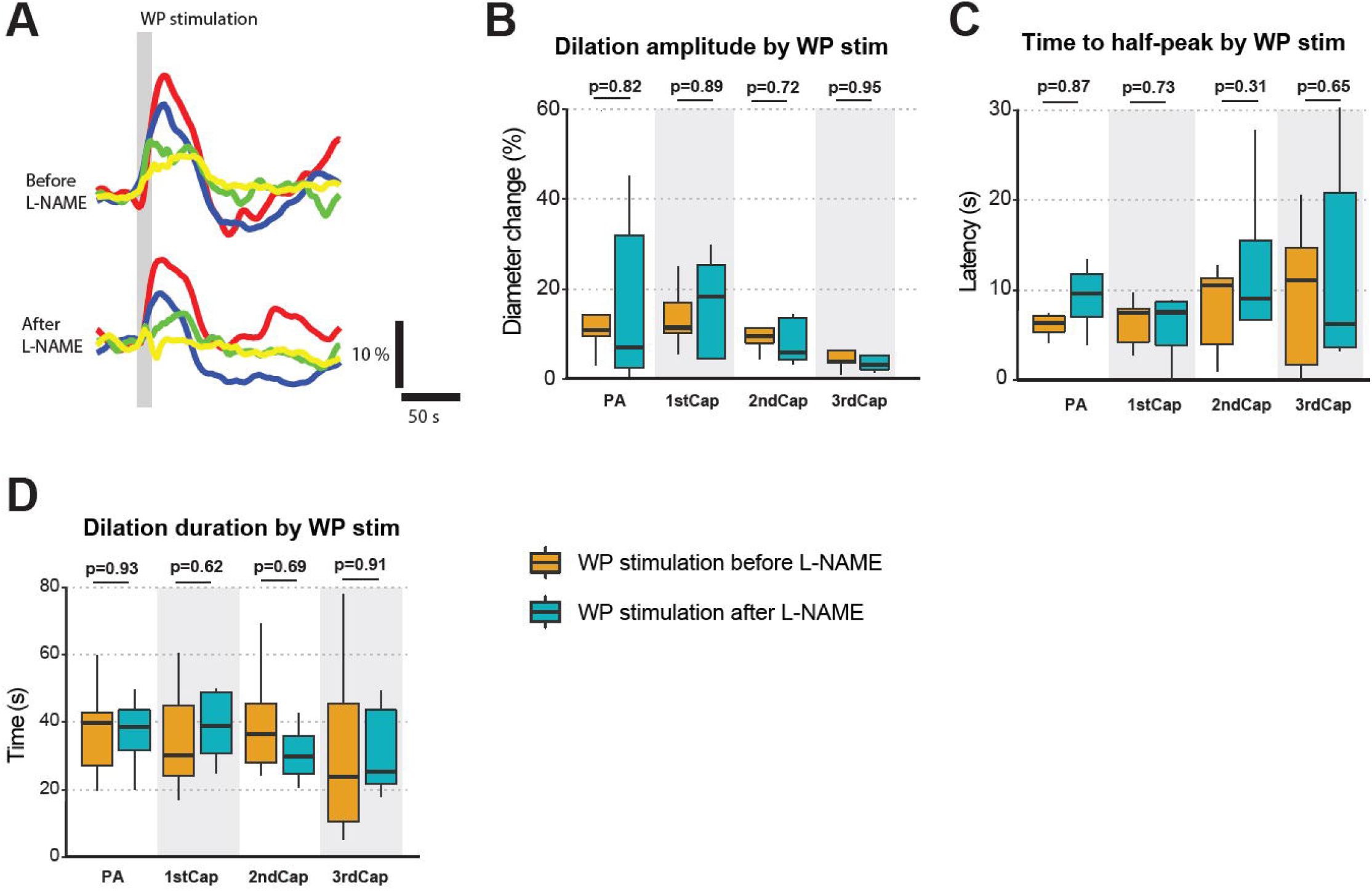
L-NAME had no significant effect on whisker pad stimulation (WP stimulation) induced vasodilation. (**A**) Representative traces of WP stimulation induced vasodilation before and after L-NAME. (**B–D**) Comparison of WP stimulation induced vasodilation before and after L-NAME, with (B) dilation amplitude, (**C**) Time to half-peak, (**D**) dilation duration of half-peak. All three measurements show no significant difference. N=5, n=5. Paired t-test.

**Supplementary figure 2:**
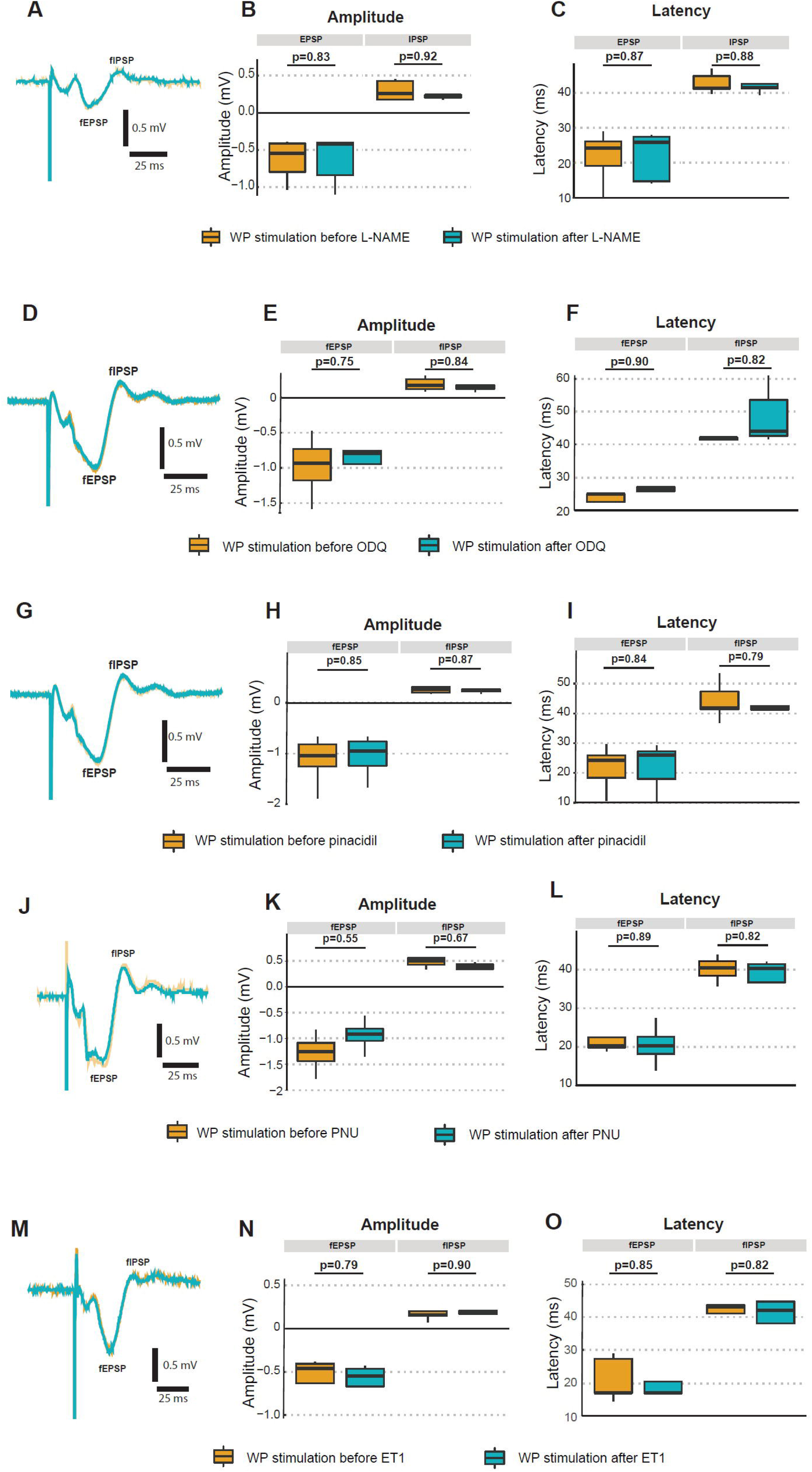
Comparison of local field potentials by WP stimulation before and after multiple vasoactive compounds. (**A–C**) Comparison of field potentials before and after i.v. bolus administration of L-NAME. (**A**) Representative traces of fEPSP and fIPSP elicited by WP stimulation before and after. (**B**) Peak amplitude of fEPSP and fIPSP. (**C**) Peak latency of fEPSP and fIPSP. (**D–F**) Comparison of field potentials before and after topical application of ODQ. (**D**) Representative traces of fEPSP and fIPSP elicited by WP stimulation before and after ODQ. (**E**) Peak amplitude of fEPSP and fIPSP. (**F**) Peak latency of fEPSP and fIPSP. Paired t-test. (**G–I**) Comparison of field potentials before and after puff of pinacidil. (**G**) Representative traces of fEPSP and fIPSP elicited by WP stimulation before and after pinacidil. (**H**) Peak amplitude of fEPSP and fIPSP. (**I**) Peak latency of fEPSP and fIPSP. (**J–L**) Comparison of field potentials before and after topical application of PNU. (**J**) Representative traces of fEPSP and fIPSP elicited by WP stimulation before and after PNU. (**K**) Peak amplitude of fEPSP and fIPSP. (**L**) Peak latency of fEPSP and fIPSP. (**M–O**) Comparison of field potentials before and after puffing of ET1. (**M**) Representative traces of fEPSP and fIPSP elicited by WP stimulation before and after ET1. (**N**) Peak amplitude of fEPSP and fIPSP. (**O**) Peak latency of fEPSP and fIPSP. Paired t-test.

**Supplementary figure 3:**
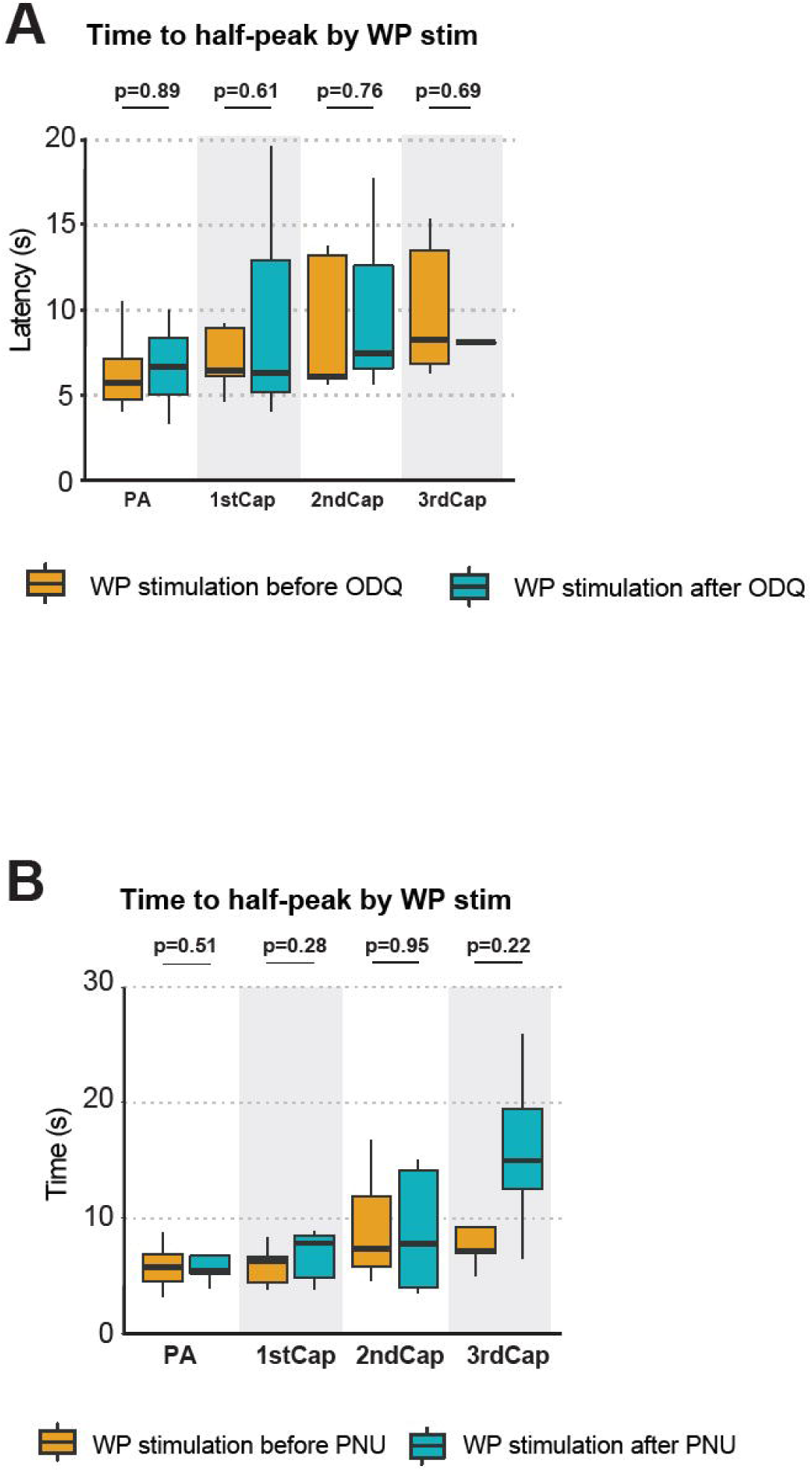
Comparison of latencies to half dilation peak by WP stimulation. (**A**) Topical application of ODQ delivered no significant change in time to half-peak for WP stimulation induced dilation. Before: N=7, n=7; after: N=6, n=6. (**B**) Topical application of PNU delivered no significant change in time to half-peak in WP stimulation induced dilation. Before: N=5, n=14; after: N=7, n=16. Unpaired t-test.

**Supplementary figure 4:**
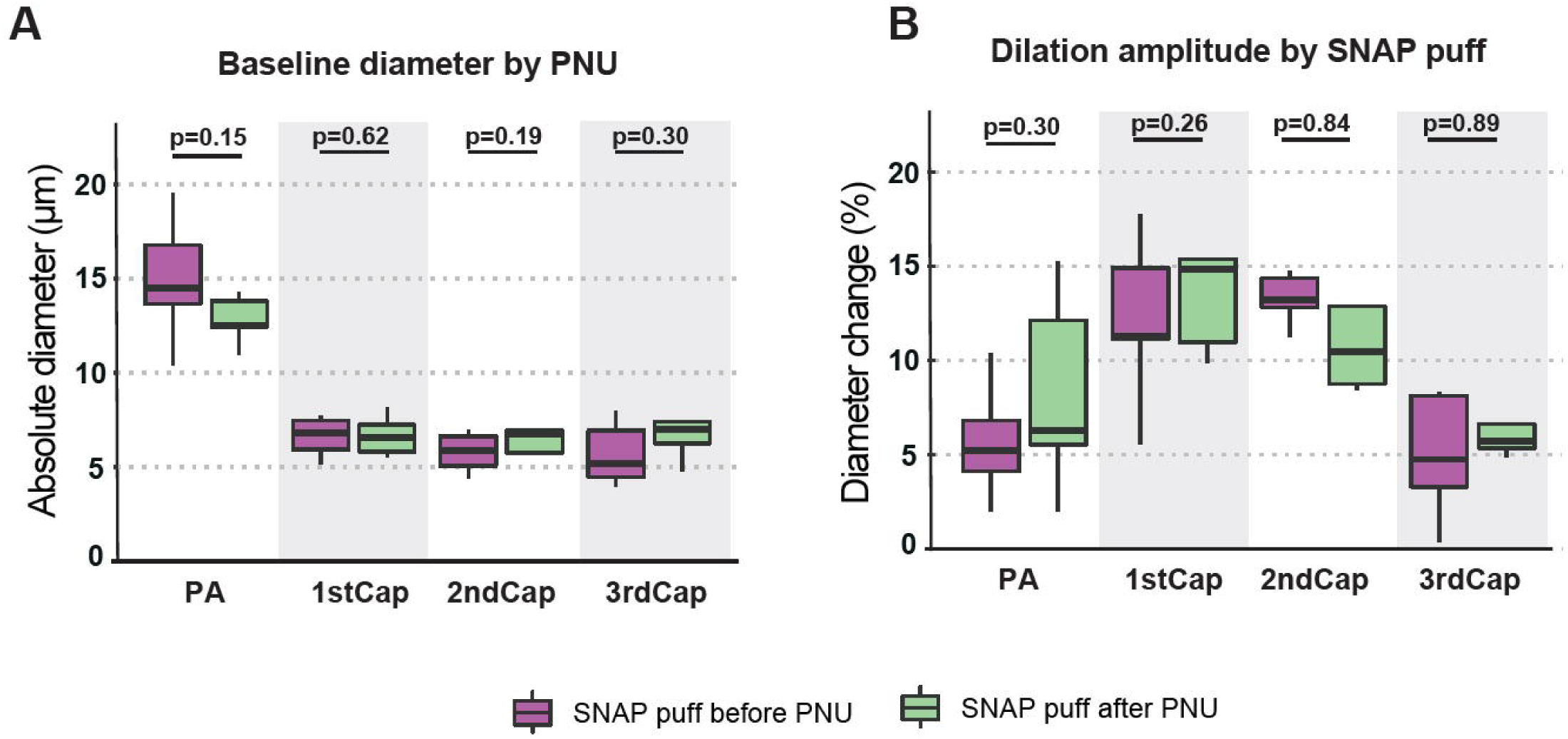
Comparison of vessel diameter change by SNAP puff before and after PNU. (**A**) Absolute vessel diameter in resting state before and after topical application of PNU. (**B**) Dilation amplitude of SNAP puff before and after topical application of PNU. Before: N=7, n=7; After: N=5, n=7. Unpaired t-test.

**Supplementary figure 5:**
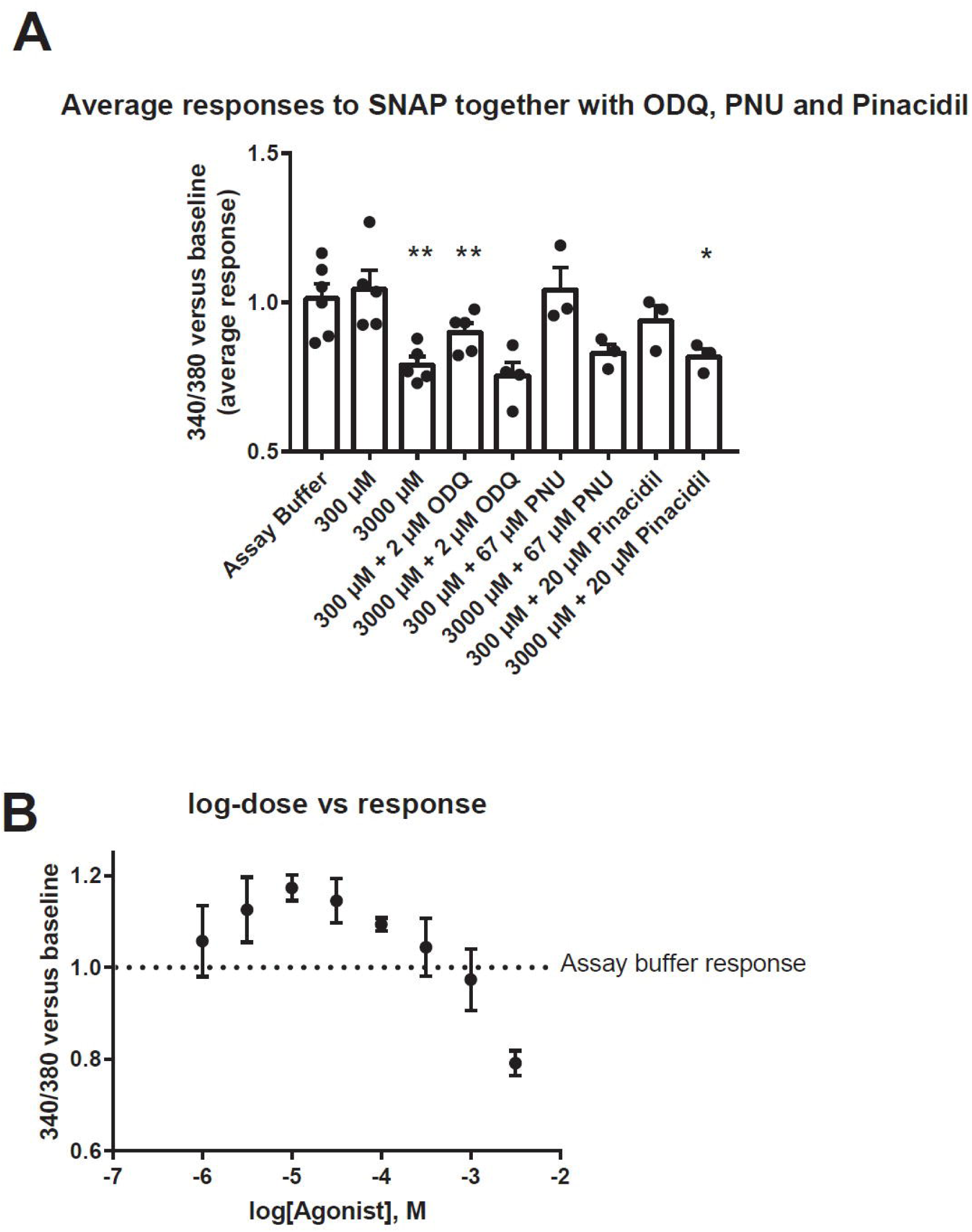
Bovine pericyte cultures respond to SNAP with a dose-dependent decrease in intracellular calcium levels. **(A)** Average responses to 300 µM or 3000 µM SNAP in emission intensity at excitation wavelengths of 340 and 380 nm. Potential inhibitors ODQ and PNU were added 15 min before the experiment, whereas pinacidil was added together with SNAP. The bars show average responses across the biological replicates (n=3–6), with the symbols indicating the average of each biological replicate. Average 340/380 ratios were compared to the response of assay buffer alone using one-way ANOVA followed by Dunnett’s multiple comparison test. (**B**) Average responses to different SNAP doses. Data points show average ± SEM of all biological replicates (n=3–6). *p<0.05, ** p<0.001, *** p<0.0001, n.s. indicates not significant.

**Supplementary figure 6.**
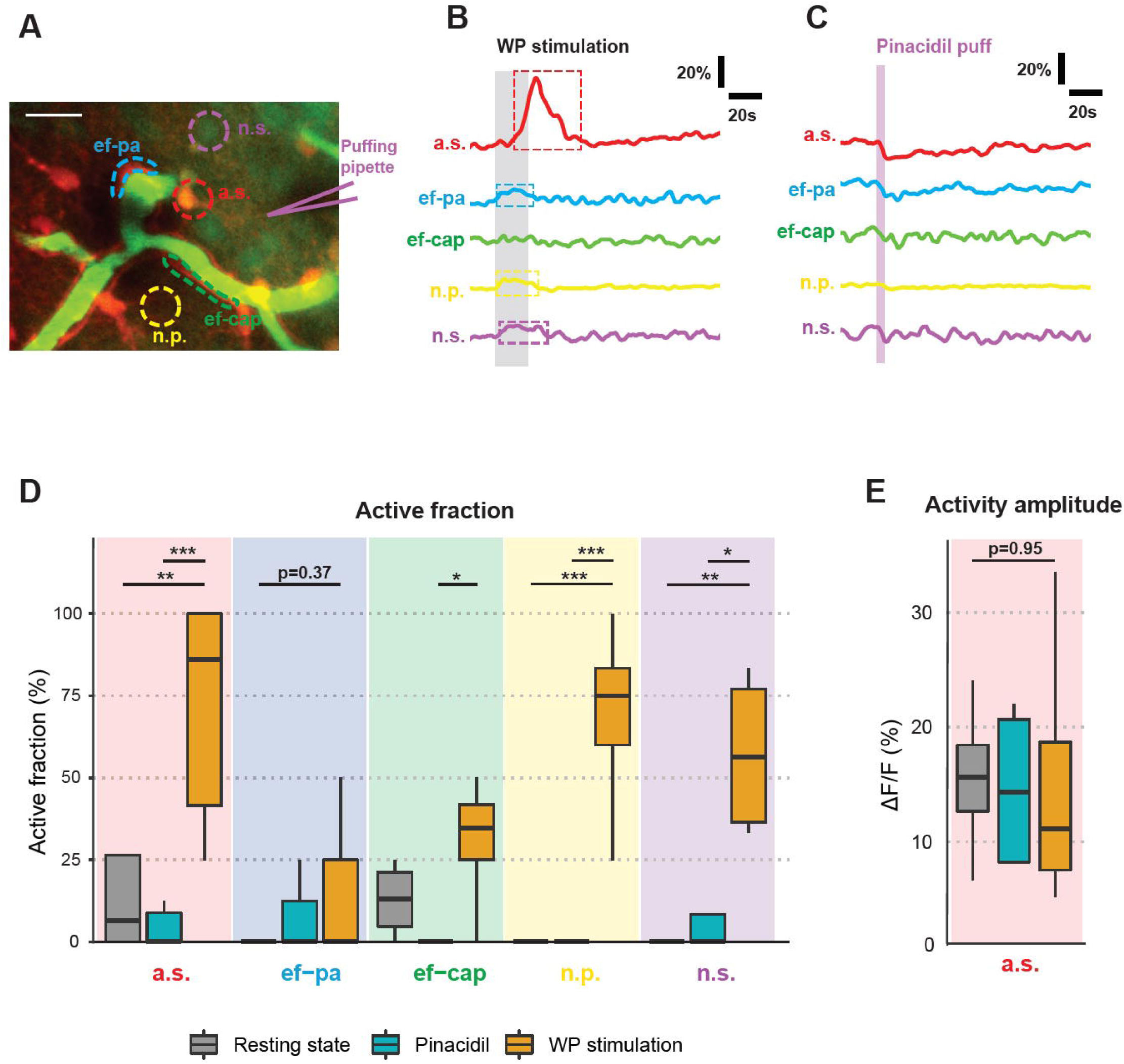
Pinacidil-induced vasodilation does not occur via neuron or astrocytes or either field potential. (**A**) The mouse brain is bulk loaded with sulforhodamin 101 (astrocyte marker as red) and Oregon Green BAPTA-1/AM (calcium indicator as green, staining both astrocytes and neurons). The two-photon image is averaged from an image z-stack. Free drawing of ROIs covering astrocytic soma (a.s.), astrocytic end-feet embracing penetrating arteriole (ef-pa), astrocytic end-feet embracing capillary (ef-cap), neuropil (n.p.), neuronal soma (n.s.). Scale bar: 10 µm. (**B**) The representative responses of each ROI upon WP stimulation. (**C**) The representative responses of each ROI upon pinacidil puff. (**D–E**) Comparison of each ROI type under three conditions: resting state, pinacidil puff and WP stimulation. (**D**) Active fraction (fraction of ROIs with at least one elicited calcium transients). (**E**) Activity amplitude (mean amplitude of all elicited calcium transients). N=10, n=149. One-way ANOVA with post hoc test.

**Supplementary figure 7.**
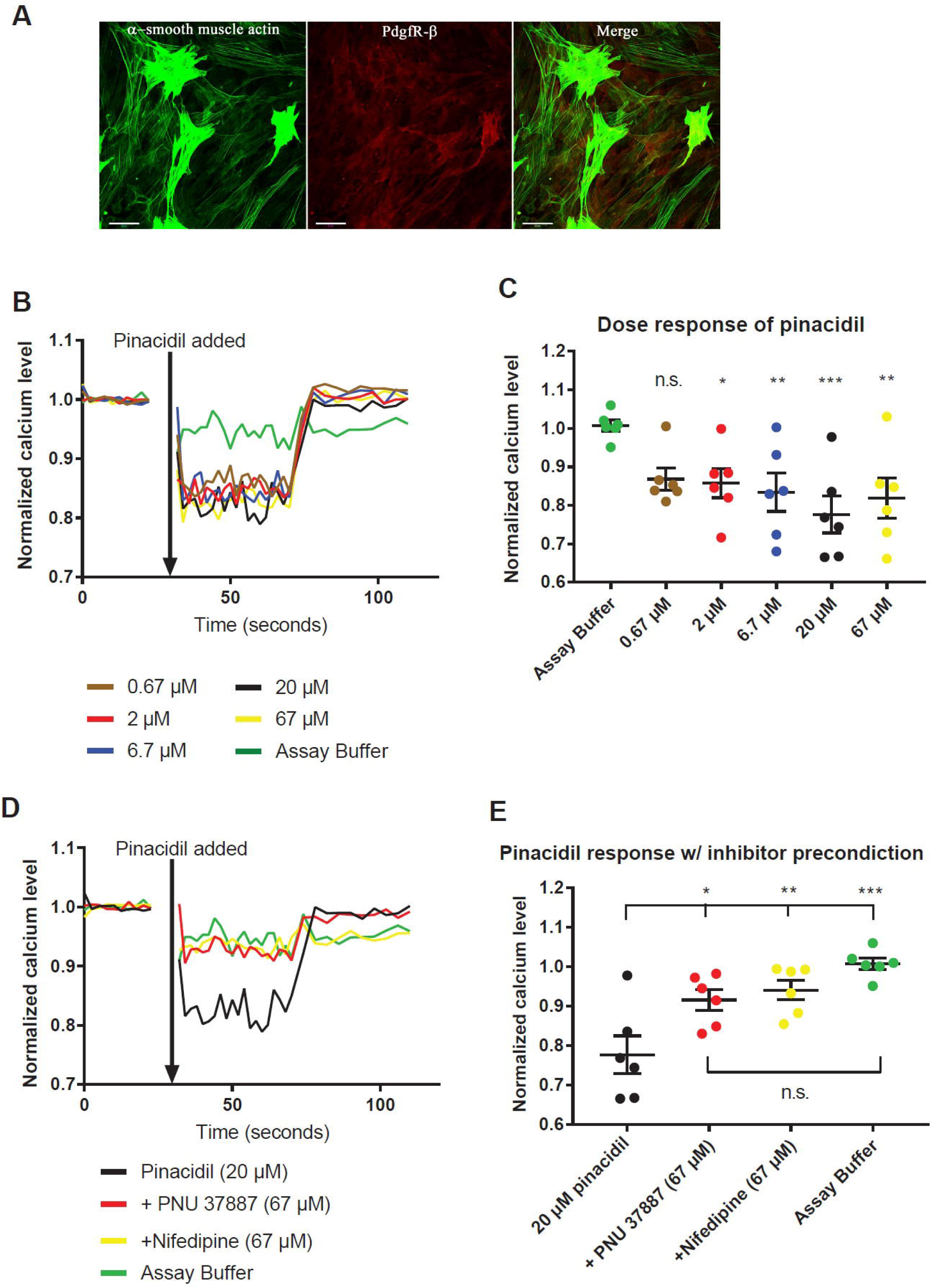
K_ATP_ channel–mediated vasodilation is via Ca^2+^-dependent pericyte signaling *in vitro*. (**A**) Immunocytochemical staining of αSMA (green), platelet-derived growth factor receptor-β (red) in monocultured bovine brain pericytes. Scale bar: 50 µm. (**B**) Representative time course of intracellular calcium curve of pericytes by adding different doses of pinacidil. (**C**) Dose-dependent responses of adding pinacidil in cultured pericytes. (**D**) Representative time course of pericyte calcium by pinacidil with or without precondition with K_ATP_ channel inhibitor 67 µM PNU or nifedipine. (**E**) Collection of 6 biological replicates confirms that both PNU and nifedipine attenuate pinacidil-induced pericyte calcium response. *p<0.05, ** p<0.001, *** p<0.0001, n.s. indicates not significant. One-way ANOVA with post hoc test.

**Supplementary figure 8:**
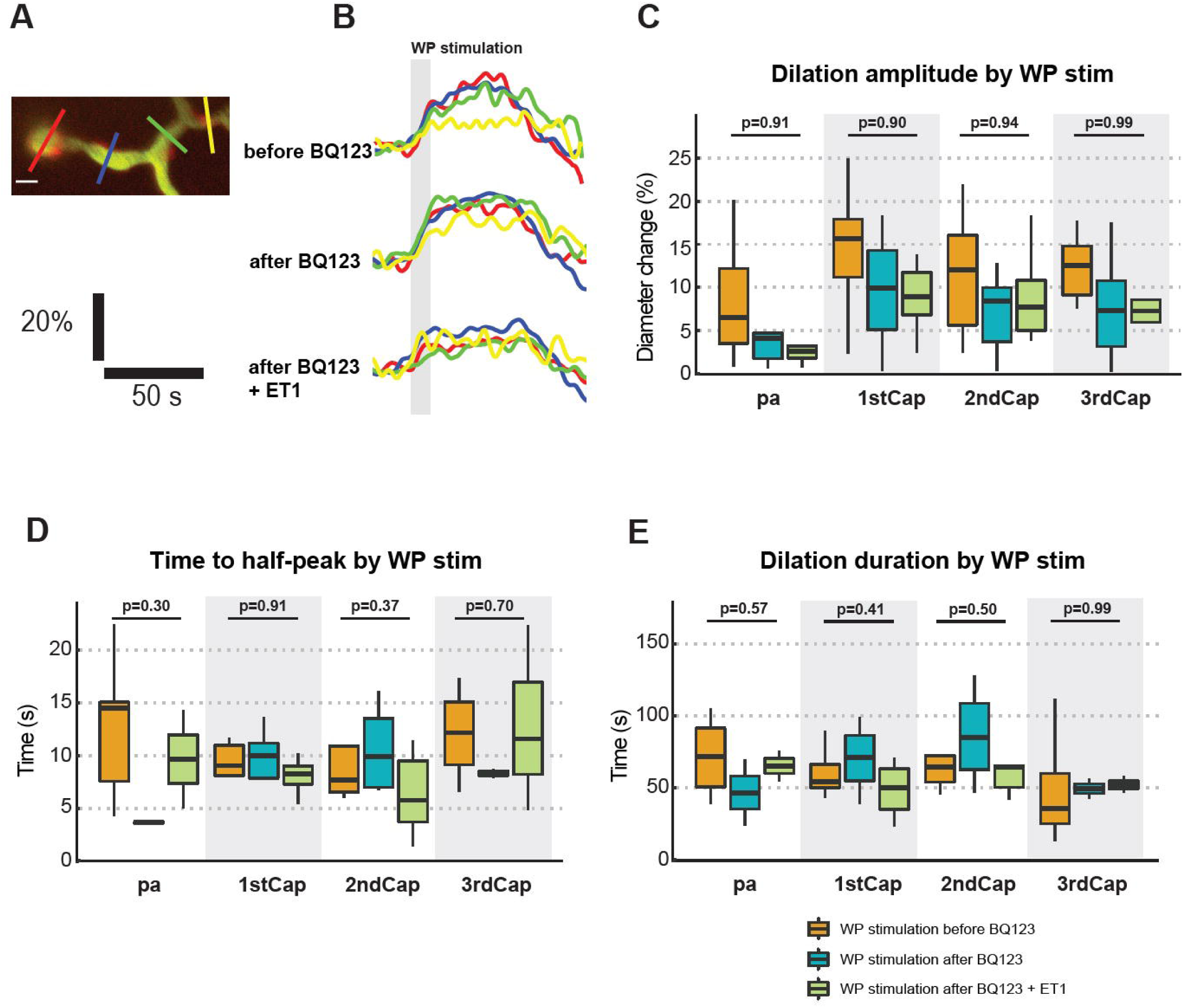
Comparison of vessel responses before BQ123 application, after BQ123 application, and after the combination of BQ123 application and ET1 puff. (**A**) An average-projected image from a local two-photon image stack with color-coded ROIs placed across PA and 1–3 order capillaries. (**B)** Representative traces of vessel dilatory curves under three conditions: before topical application of BQ123, after topical application of BQ123, and after the combination of BQ123 application and ET1 puff. (**C**) Comparison of dilation amplitude by WP stimulation. (**D**) Comparison of time to half-peak. (**E**) Comparison of half-peak dilation duration. All three conditions: N=6, n=11. One-way ANOVA with post-hoc test.

**Supplementary figure 9:**
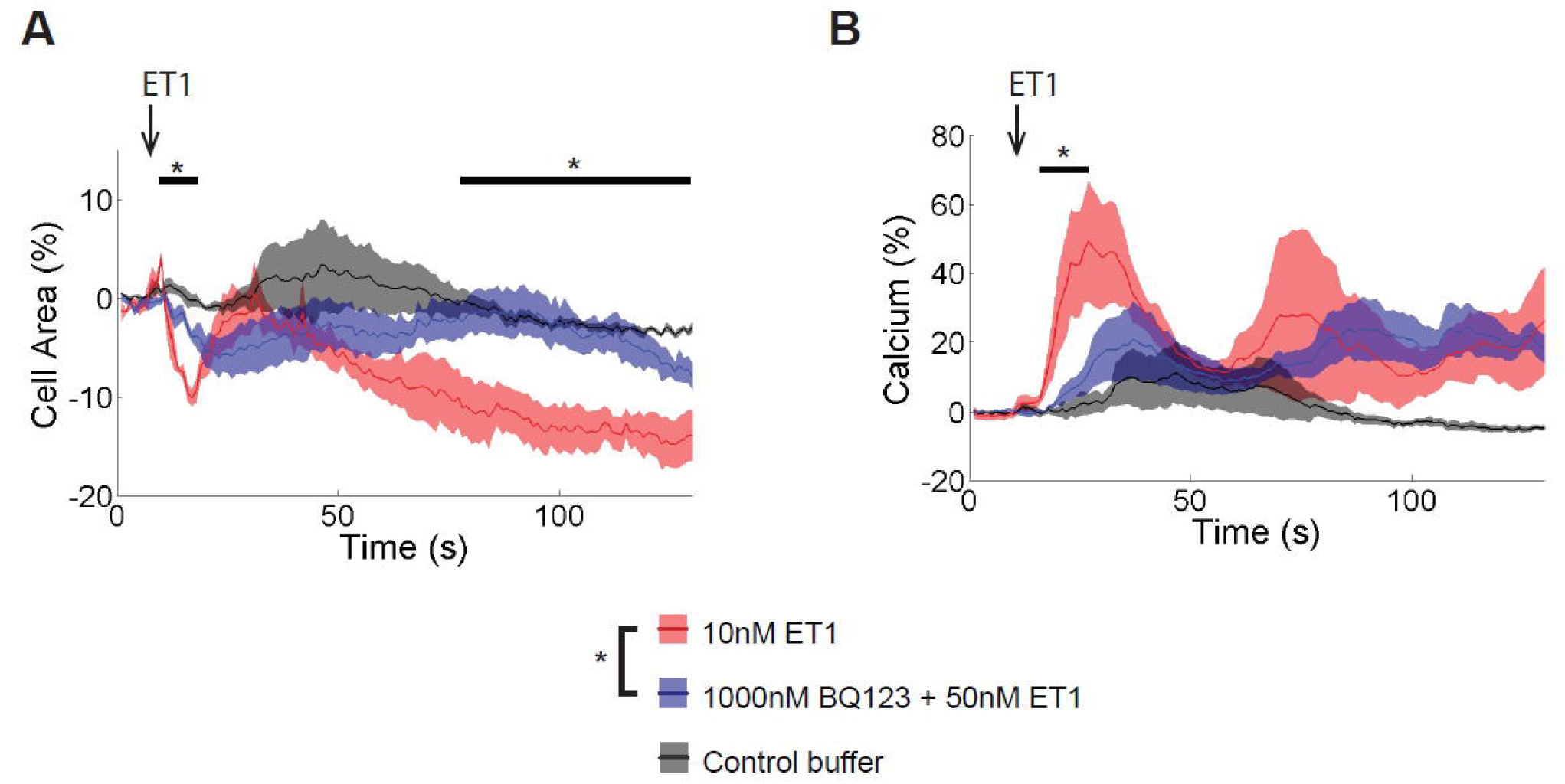
Contraction and calcium rise in cultured bovine pericytes by the administration of ET-1 and BQ123. (**A**) Pericyte area change upon administration of ET1 with or without precondition with 1000 nM BQ123. (**B**) Comparison of pericyte calcium increase. We chose to compare 10 nM ET1 with 50 nM ET1 under BQ123. This was because 50 nM ET1 induced large calcium rise and therefore false increase in cell area by our analysis algorithm. 10 nM ET1 triggered larger pericyte constriction than 50 nM ET1 under BQ123, proving the specificity of vasoconstricting effect of ET1 channels. Unpaired t-test, *p<0.05. N=4, n=355.

**Supplementary figure 10.**
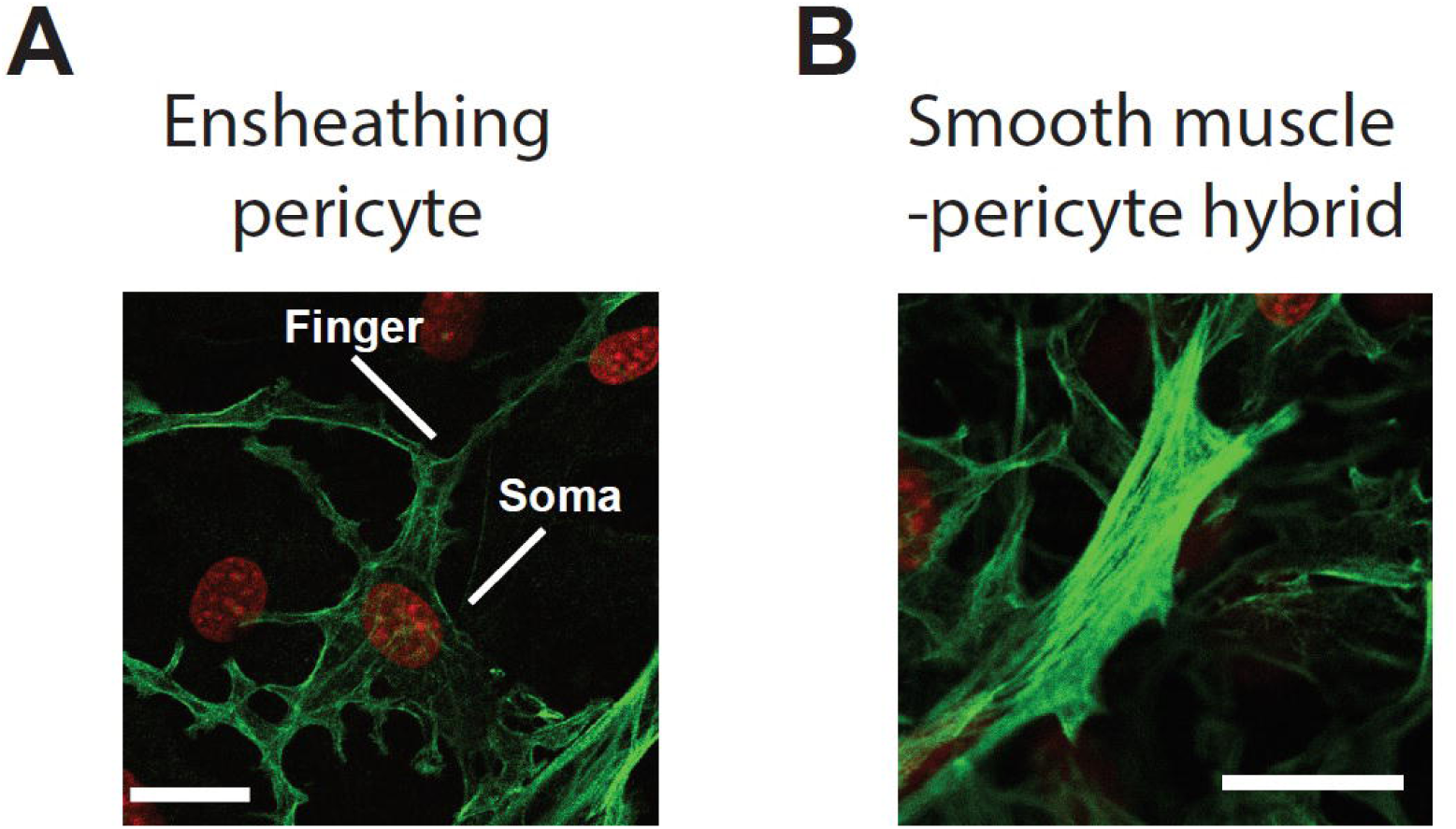
Representative immunohistochemical staining in culture bovine pericytes. (**A–B**) Immunoohistochemistry stains αSMA as green and neucleus and red. αSMA marks (**A**) somas and fingers of ensheathing pericytes, and (**B**) somas of smooth muscle pericyte hybrids. Scale bar: 20 µm.

**Supplementary figure 11:**
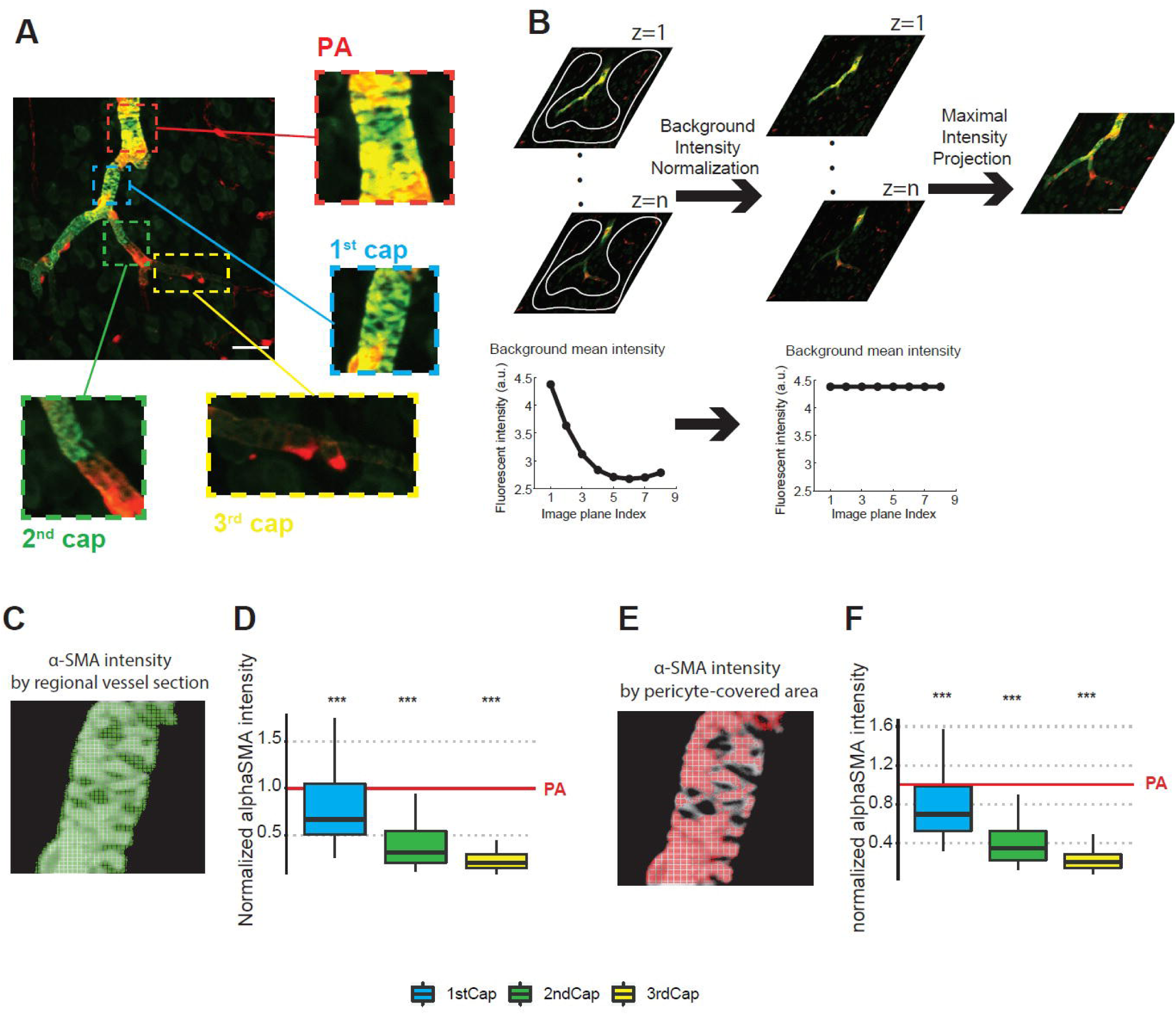
Quantification of αSMA density at PA, 1–3 order capillaries. (**A**) Immunohistochemistry stains αSMA as green, and pericytes and smooth muscle cells are endogeneously stained as red. The image is a maximal intensity project from an image z-stack. Different vessel segments are identified as PA and 1–3 order capillaries. (**B**) The image processing procedure for the z-stack is to free-draw background area and then calculate the mean background fluorescent intensity for each image plane. The background intensity is normalized to the peak value of the z-stack. Finally, the z-stack images are projected onto one image by maximal projection. (**C–D**) Regarding the measurement of αSMA intensity, two methods are used. (**C**) Hand-drawn delineation of the vessel region in the green image, and calculation of the mean intensity in the selected region. (**D**) The red image is used to select the pericyte-positive pixels and to calculate the mean intensity of the corresponding pixels in the green image. (**E–F**) For each z-stack image, the αSMA intensity is normalized to PA. 1–3 order capillaries show significantly smaller αSMA intensity compared to PA by both methods, using method (**C**) for plot (**D**), and method (**E**) for plot (**F**). N=3, n=31. One-way ANOVA with post hoc test.

