## Supplementary material for "Nitric oxide, K_ATP_ channels and endothelin-1 modulate brain pericyte function, vascular tone and neurovascular coupling"

**Pinacidil does not impact neuronal or astrocytic [Ca^2+^]_i_, neither elicited field potentials**

K_ATP_ channels are abundant in brain pericytes [1, 2], but K_ATP_ channels also prevail in neurons and astrocytes [3, 4]. Therefore, in this type of study one must consider the possibility that the effect of pinacidil on microvascular responses reflected an effect on underlying neurons or glia rather than on smooth muscle cells and pericytes. To assess this possibility, we examined the neuronal and astrocytic Ca^2+^ responses in response to pinacidil puffing on blood vessels.

Regions of interest (ROIs) were placed to study different compartments of astrocytes, neurons and neuropil near PA and 1-3 order capillaries (**Supplementary Fig. 4A**). WP stimulation induced intracellular Ca^2+^ transient (see definition in the Methods) in astrocytic soma, astrocytic endfeet embracing the PA, neuropil, and neuronal soma (**Supplementary Fig. 4B**). Puffing of pinacidil at the same location did not evoke Ca^2+^ transients in any of the subcellular compartments of nerve cells (**Supplementary Fig. 4C and 4D**), consistent with a previous in vitro study [5]. Likewise, the amplitude of astrocytic soma with a Ca^2+^ transient was the same in the resting state as during pinacidil puff and WP stimulation (**Supplementary Fig. 4E**).
